# Identification of evolutionarily conserved virulence factor by selective pressure analysis of *Streptococcus pneumoniae*

**DOI:** 10.1101/356212

**Authors:** Masaya Yamaguchi, Kana Goto, Yujiro Hirose, Yuka Yamaguchi, Tomoko Sumitomo, Masanobu Nakata, Kazuhiko Nakano, Shigetada Kawabata

## Abstract

Evolutionarily conserved virulence factors can be candidate therapeutic targets or vaccine antigens. Here, we investigated the evolutionary selective pressures on 16 pneumococcal choline-binding cell-surface proteins since *Streptococcus pneumoniae* is one of the pathogen posing the greatest threats to human health. Phylogenetic and molecular analyses revealed that *cbpJ* had the highest codon rates to total numbers of codons under significant negative selection among those examined. Our *in vitro* and *in vivo* assays indicated that CbpJ functions as a virulence factor in pneumococcal pneumonia by contributing to evasion of neutrophil killing. Deficiency of *cbpL* under relaxed selective pressure also caused a similar tendency but showed no significant difference in mouse intranasal infection. Thus, molecular evolutionary analysis is a powerful tool that reveals the importance of virulence factors in real-world infection and transmission, since calculations are performed based on bacterial genome diversity following transmission of infection in an uncontrolled population.

---

Improper use of antibiotics creates evolutionary pressures that drive bacteria to acquire drug resistance by natural mutation and/or horizontal transfer of resistance genes. This is a major public health threat: it is estimated that drug-resistant infections cause 10 million deaths annually and may result in economic losses reaching 100 trillion US dollars by 2050^1^. However, a target-to-hit screen typically requires approximately 24 discovery projects and 94 million US dollars, and the baseline total cost is 1.8 billion US dollars over 13 years to launch a new drug^2^. In fact, the number of new antibiotics developed and approved has steadily decreased in the past three decades, leaving fewer options for treating resistant bacteria^3^.

*Streptococcus pneumoniae* is one of the pathogens posing the greatest threat to human health^4,5^. *S. pneumoniae* belongs to the mitis group^6,7^ and is a major cause of pneumonia, sepsis, and meningitis^8,9^. In 2015, pneumococcal pneumonia caused over 1.5 million deaths in individuals of all ages, and this rate increased in people over 70 years old between 2005 and 2015^10^, which is especially problematic since the elderly population is growing in many parts of the world. Although pneumococcal conjugate vaccines have considerable benefits, non-vaccine pneumococcus serotypes have increased worldwide^11,12^.

Conflict between the host immune system and pathogens leads to an evolutionary arms races known as the “Red Queen” scenario^13,14^. Protein regions at the host–pathogen interface are subjected to the strongest selective pressure and thus evolve under positive selection. Adaptive evolution has been reported in genes related to the mammalian immune system such as pattern recognition receptors^14^. Concerning negative/purifying selection, Jordan *et al.* compared two whole genome sequences and showed that essential bacterial genes appear to demonstrate substantially lower average values of synonymous and nonsynonymous nucleotide substitution rates compared to those in nonessential genes^15^. However, to our knowledge, comprehensive evolutional analysis on codons of genes encoding bacterial cell surface proteins has not been performed. Mutations on essential genes directly cause host death because essential genes encode proteins to maintain basic bacterial survival such as central metabolism, DNA replication, translation of genes into proteins, and so on. Meanwhile, nonessential genes are under negative/purifying selection, which is important for the survival and/or success of the species in the host and/or the environment as non-synonymous substitution of codons can lead to lineage extinction (Fig. 1). Phylogenetic and molecular evolutionary analyses can reveal the number of codons under negative/purifying selection in a species. Because alterations in amino acid residues in regions under negative selective pressure are not allowed, drugs targeting these regions would be less likely to promote the development of resistance through natural mutation.

**Figure 1.**
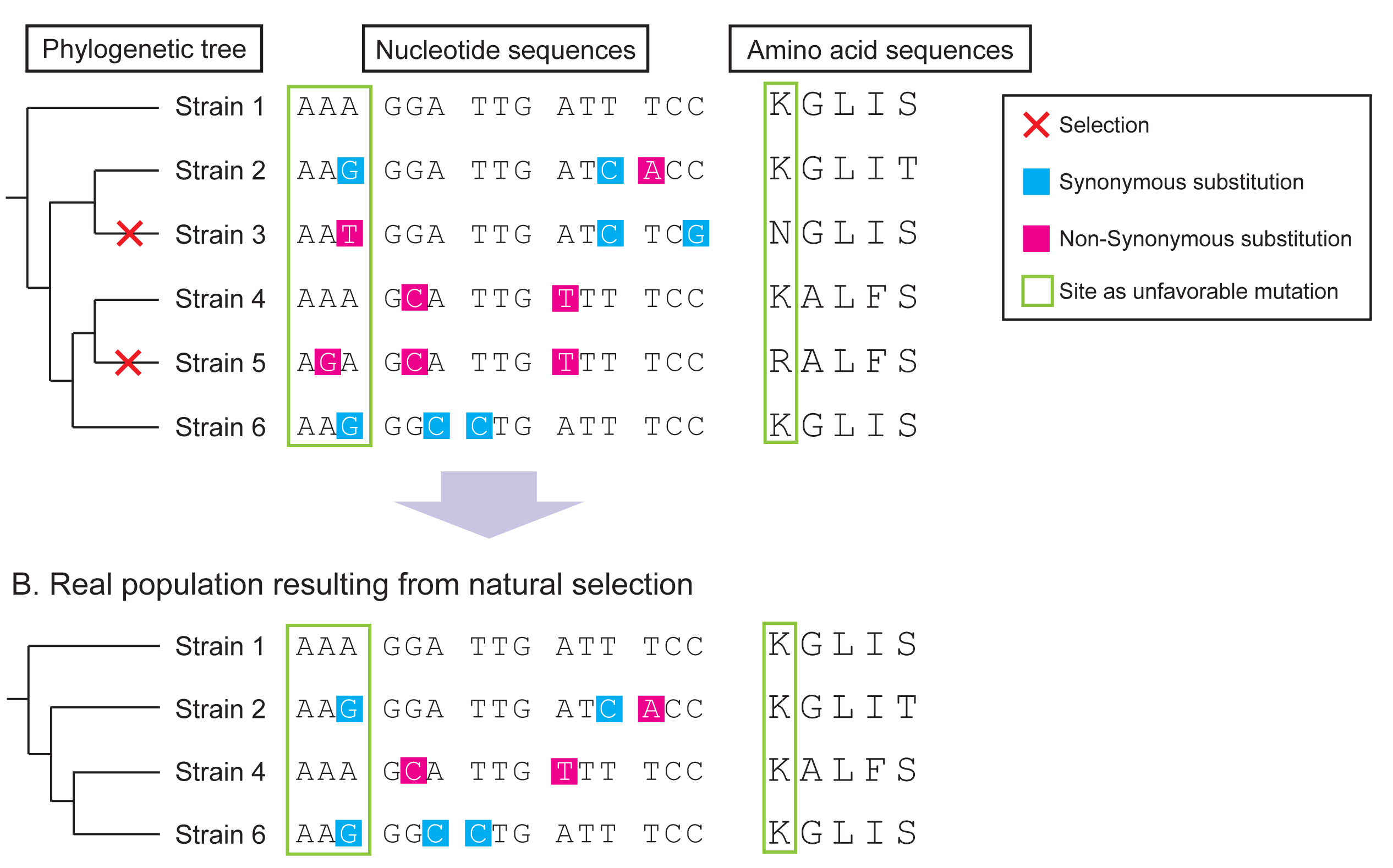
Scheme for intra-species molecular evolutionary analysis. **A.** Random genetic drift induces synonymous and non-synonymous mutations with equal probability. However, non-synonymous mutations in the essential region cause host selection. **B.** As a result of natural selection, synonymous substitutions are concentrated in important genes. Phylogenetic and molecular evolutionary analyses can detect significant accumulation of synonymous substitutions in codons of host proteins. Codon-based analysis yields much more information than nucleotide- or amino acid-based analyses.

We analysed pneumococcal choline-binding proteins (CBPs) localised on the bacterial cell surface through interaction with choline-binding repeats and phosphoryl choline on the cell wall. At least some CBPs play key roles in cell wall physiology, in pneumococcal adhesion and invasion, and in evasion of host immunity. *S. pneumoniae* harbours various CBPs including *N*-acetylmuramoyl L-alanine amidase (LytA), which induces pneumococcal-specific autolysis^16-18^. Pneumococcal surface protein A (PspA) is a highly variable protein and inhibits complement activation^17-20^. Choline binding protein A (CbpA; also called PspC) works as a major pneumococcal adhesin and contributes to evasion of host immunity via interaction with several host proteins^17,18,21^. Choline binding protein L (CbpL) contains the choline binding repeats sandwiched between the Excalibur and lipoproteins domains and works as an anti-phagocytic factor^22^. Although several CBPs have been characterised, their phylogenetic relationships remain unclear and the unclassified gene names are confusing. We first analysed the distribution of genes encoding CBPs based on pneumococcal genome sequences. Orthologues of genes in each strain were identified by phylogenetic analysis. We then calculated the evolutionary selective pressure on each codon from the phylogenetic trees and aligned sequences. We found that *cbpJ* contains the highest rate of codons under negative selection. CbpJ has no known functional domains except signal sequences and choline-binding repeats, and its role in pneumococcal pathogenesis is unclear. Functional analyses revealed that CbpJ contributes to evasion of host neutrophil-mediated killing in pneumococcal pneumonia. Thus, evolutionary analysis focusing on negative selection can reveal novel virulence factors.

## Results

### Distribution of *cbp* genes among pneumococcal strains

Genes encoding CBPs among pneumococcal strains were extracted by tBLASTn search (Supplementary Table 1). Some genes were re-annotated since the search results showed that certain homologous regions were not matched to annotated open reading frames (ORFs). In strain SPNA45, *SPNA_01670* contains both predicted promoter regions and intact ORF structures of *cbpF* and *cbpJ*. On the other hand, *cbpG*-homologous regions in strains R6, D39, SPN034183, SPN994038, and SPN994039 did not contain promoters (Supplementary Table 1 and Supplementary Table 2). Orthologous relationships of each gene were analysed. The distribution of *cbp* genes was not correspondent with capsular serotypes (Fig. 2A). Four genes—i.e., *lytA, lytB, cbpD*, and *cbpE*—were conserved as intact ORFs in all 28 pneumococcal strains (Fig. 2A). Other *cbp* genes contained frameshift mutations in the orthologues or were absent in some strains.

**Figure 2.**
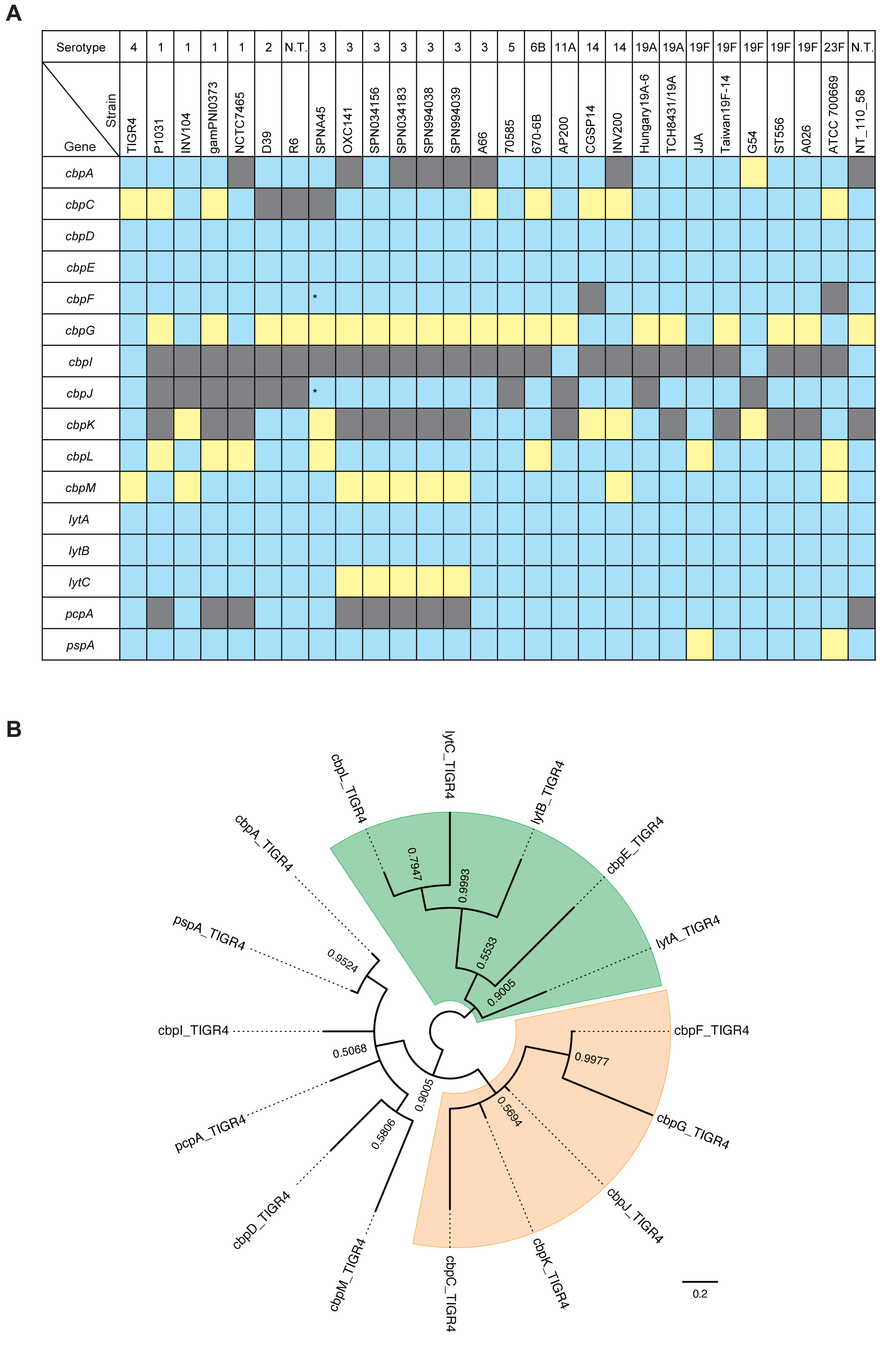
Distribution of *cbp* genes and phylogenetic relationship in TIGR4. **A.** Distribution of genes encoding CBPs among pneumococcal strains. The gene locus tag numbers are shown in Supplementary Table 1. Blue, yellow, and gray show the presence, pseudogenisation, and absence of genes, respectively. *These genes are annotated as one gene, but our bioinformatic analysis indicates that they are independent genes. **B.** Nucleotide-based Bayesian phylogenetic tree of *cbp* genes of *S. pneumoniae* strain TIGR4. The tree is unrooted and posterior probabilities are shown near the nodes. The scale bar indicates nucleotide substitutions per site.

### Phylogenetic relationships in pneumococcal CBPs

Phylogenetic relationships of genes encoding CBPs in pneumococcal species are confusing since some genes in the same cluster show high similarity to each other. To clarify the relationships, we compared common nucleotide sequences among genes encoding CBPs in the strain TIGR4. Maximum likelihood and Bayesian phylogenetic analyses revealed two common clusters: one comprising *cbpF, cbpG, cbpJ, cbpK*, and *cbpC*, and the other comprising *lytA, lytB, lytC, cbpL*, and *cbpE* (Fig. 2B and Supplementary Fig. 1). The names of some CBP genes were not consistent with those of phylogenetically related genes. In particular, *cbpF, cbpG, cbpJ*, and *cbpK* were located close to each other in pneumococcal genomes and showed high similarity. We thus defined orthologous genes in each pneumococcal strain based on maximum likelihood and Bayesian phylogenetic analyses (Fig. 3 and Supplementary Fig. 2). The gene locus tag numbers in orthologous relationships are shown in Supplementary Table 1. The sequence similarity of *cbpF, cbpG, cbpJ*, and *cbpK* and their close proximity within genomes indicated that a common ancestral *S. pneumoniae* acquired the genes by duplication. Phylogenetic trees showed well-separated clusters of each gene. These independent relationships indicated that horizontal gene transfer did not contribute to the spread of *cbpF, cbpG, cbpJ*, and *cbpK* in *S. pneumoniae* species, despite their ability to take up exogenous DNA. The genetic diversity of these genes may have been established by accumulation of natural mutations during pneumococcal transmission.

**Figure 3:**
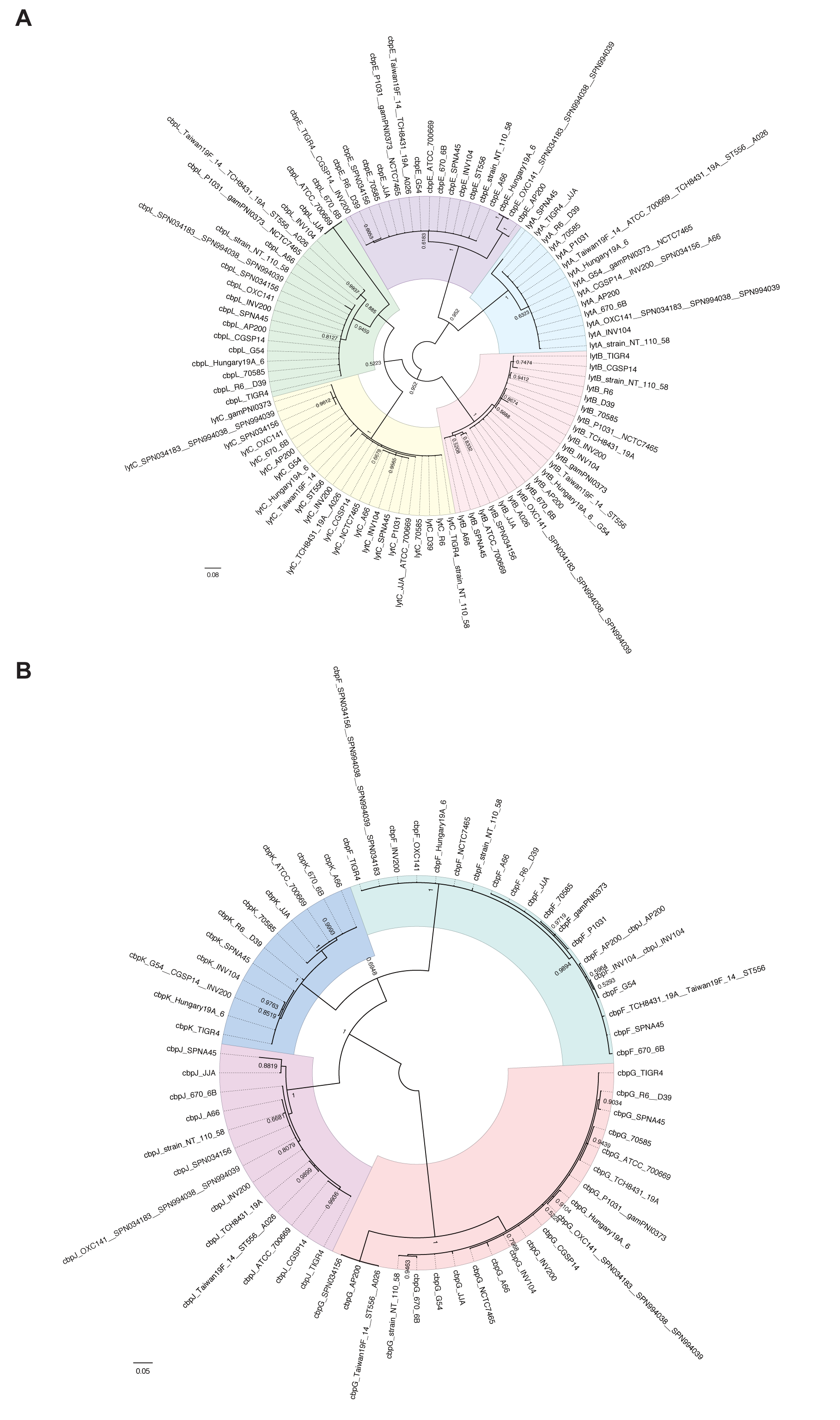
Phylogenetic analyses of *cbp* genes with high similarity. **A, B.** Nucleotide-based Bayesian phylogenetic tree of the *lytA, lytB, lytC, cbpE*, and *cbpL* genes (A) and the *cbpF, cbpG, cbpJ*, and *cbpK* genes (B) in *S. pneumoniae*. The trees are unrooted although they are presented as midpoint-rooted for clarity. Strains with identical sequences are listed on the same branch. Posterior probabilities are shown near the nodes. The scale bar indicates nucleotide substitutions per site.

### Evolutionary selective pressures on each of the CBP codons

To evaluate the significance of CBPs in real-life infection and transmission, we performed molecular evolutionary calculations based on bacterial genome diversity established after transmission of infection in an uncontrolled population. The nucleotide sequences of each CBP were aligned by codon, and conserved common codons were used for phylogenetic analysis (Supplementary Fig. 3). The selective pressure on each gene was calculated based on the phylogenetic trees and aligned sequences (Table 1). The rates of codons under negative selection are visualised in Supplementary Figure 4. Over 13% of total codons in *cbpJ* and *lytA* were under negative selection compared to less than 5% for other *cbp* genes, indicating that these genes play an important role in the success of *S. pneumoniae* species. On the other hand, *pspA* encoding the genetically divergent virulence factor PspA, contained fewer evolutionarily conserved codons, but had the highest numbers of codons under positive pressure. Additionally, there were no evolutionarily conserved codons in *cbpG, cbpC*, and *cbpL*. The latter two had no common codons as few genes had frameshift mutations. When we re-calculated selective pressure without these genes, we found a low rate of codons under negative selection among CBP-encoding genes (Supplementary Table 3).

**Table 1.** Evolutionary analyses of genes encoding choline-binding proteins*

| Genes | Number of sequences<sup>1</sup> | dN/dS | Coverage of comparable codons relative to whole protein in TIGR4 | Codons evolving under positive selection | Codons evolving under purifying selection | % Of codons under purifying selection relative to total codons |
| --- | --- | --- | --- | --- | --- | --- |
| *cbpA* | 19 | 0.864 | 22.334% (155/694) | 3.226% (5/155) | 7.742% (12/155) | 1.729% |
| *cbpC* | 13 | – | 0% (0/93) | – | – | 0.000% |
| *cbpD* | 19 | 0.359 | 75.278% (338/449) | 0.296% (1/338) | 3.550% (12/338) | 2.672% |
| *cbpE* | 18 | 0.325 | 99.363% (624/628) | 0.160% (1/624) | 4.968% (31/624) | 4.936% |
| *cbpF* | 19 | 0.395 | 60.411% (206/341) | 0.485% (1/206) | 3.398% (7/206) | 2.053% |
| *cbpG* | 21 | – | 0% (0/286) | – | – | 0.000% |
| *cbpI* | 2 | – | – | – | – | – |
| *cbpJ* | 15 | 0.346 | 84.084% (280/333) | 1.429% (4/280) | 18.571% (52/280) | 15.616% |
| *cbpK* | 11 | 0.353 | 85.630% (292/341) | 0.342% (1/292) | 3.082% (9/292) | 2.639% |
| *cbpL* | 20 | – | 0% (0/333) | – | – | 0.000% |
| *cbpM* | 10 | 0.642 | 98.462% (128/130)<sup>2</sup> | 0% (0/128) | 0% (0/128) | 0.000% |
| *lytA* | 14 | 0.141 | 80.564% (257/319) | 0% (0/257) | 17.121% (44/257) | 13.793% |
| *lytB* | 22 | 0.185 | 92.868% (612/659) | 0% (0/612) | 4.739% (29/612) | 4.401% |
| *lytC* | 23 | 0.400 | 19.348% (95/491) | 0% (0/95) | 5.263% (5/95) | 1.018% |
| *pcpA* | 18 | 0.261 | 77.010% (479/622) | 0% (0/479) | 0.418% (2/479) | 0.322% |
| *pspA* | 24 | 0.857 | 19.060% (142/745) | 6.338% (9/142) | 12.676% (18/142) | 2.416% |

<sup>1</sup>Sequences with 100% identity were treated as the same sequence; <sup>2</sup>compared to D39.
\*Evolutionary analysis was performed by Bayesian inference of aligned *cbp* sequences from complete genomes of *S. pneumoniae* with the two-rate fixed-effects likelihood function in HyPhy software package. dN/dS is the ratio of non-synonymous to synonymous changes in overall analysed genes. Individual codons with a statistically significant signature were also calculated and are expressed as a percentage of the total number of codons included in the analysis.

### CbpJ acts as a virulence factor in pneumococcal pneumonia

While CbpJ had the highest rate of codons under negative selection among pneumococcal CBPs, it has no known functional domains except a choline-binding repeat in its amino acid sequence. Moreover, its role in pneumococcal pathogenesis is unknown. In contrast, CbpL had no common comparable codons and showed limited numbers of evolutionarily conserved codons even after the above-described adjustment. The domain structures and codons of CbpJ and CbpL under negative selection are shown in Figure 4A. The domains were searched using MOTIF Libraries including PROSITE, NCBI-CDD, and P-fam^23-26^. To assess the roles of CbpJ and CbpL in pneumococcal pathogenesis, we generated mutant strains deficient in the corresponding genes. The mutant strains showed a slightly steeper growth curve in THY medium (Supplementary Fig. 5A). There were no differences among the strains in minimum inhibitory concentration (MIC) and minimum bactericidal concentration (MBC) values for penicillin G, and bacterial morphology (Supplementary Table 4 and Supplementary Fig. 5B). WT and mutant strains in stationary phase showed that most cells were stained violet, whereas almost all cells of strains in the decline phase were stained pink probably due to autolysis (Supplementary Fig. 5B). The *lytA* gene expression was slightly increased in the *ΔcbpJ* strain compared to that in the WT strain at the log and decline phases (Supplementary Fig. 5C). However, as described above, the difference did not seem to affect pneumococcal autolysis substantially. We first performed a mouse intranasal infection assay to investigate the role of CbpJ and CbpL in pneumonia. Mice intranasally infected with strain *ΔcbpJ* showed an improved survival rate compared to those infected with WT *S. pneumoniae*; although a similar tendency was observed for *ΔcbpL*-infected relative to WT mice; the difference was not statistically significant (Fig. 4B). The number of bacteria in the bronchoalveolar lavage fluid (BALF) from *ΔcbpJ*-infected mice was lower than that in the BALF from *ΔcbpL*- and WT-infected mice (Fig. 4C). We also performed competitive assay by intranasal co-infection with the WT and *ΔcbpJ* strains. The BALF at 24 h after infection showed fewer bacterial CFUs of *ΔcbpJ* compared to those of the WT (Fig. 4D). We also examined whether CbpL or CbpJ contributes to the association of *S. pneumoniae* with alveolar epithelial cells and found that WT *S. pneumoniae* as well as *ΔcbpL* and *ΔcbpJ* mutant strains did not differ in their ability to adhere to A549 human alveolar epithelial cells (Fig. 4E).

**Figure 4:**
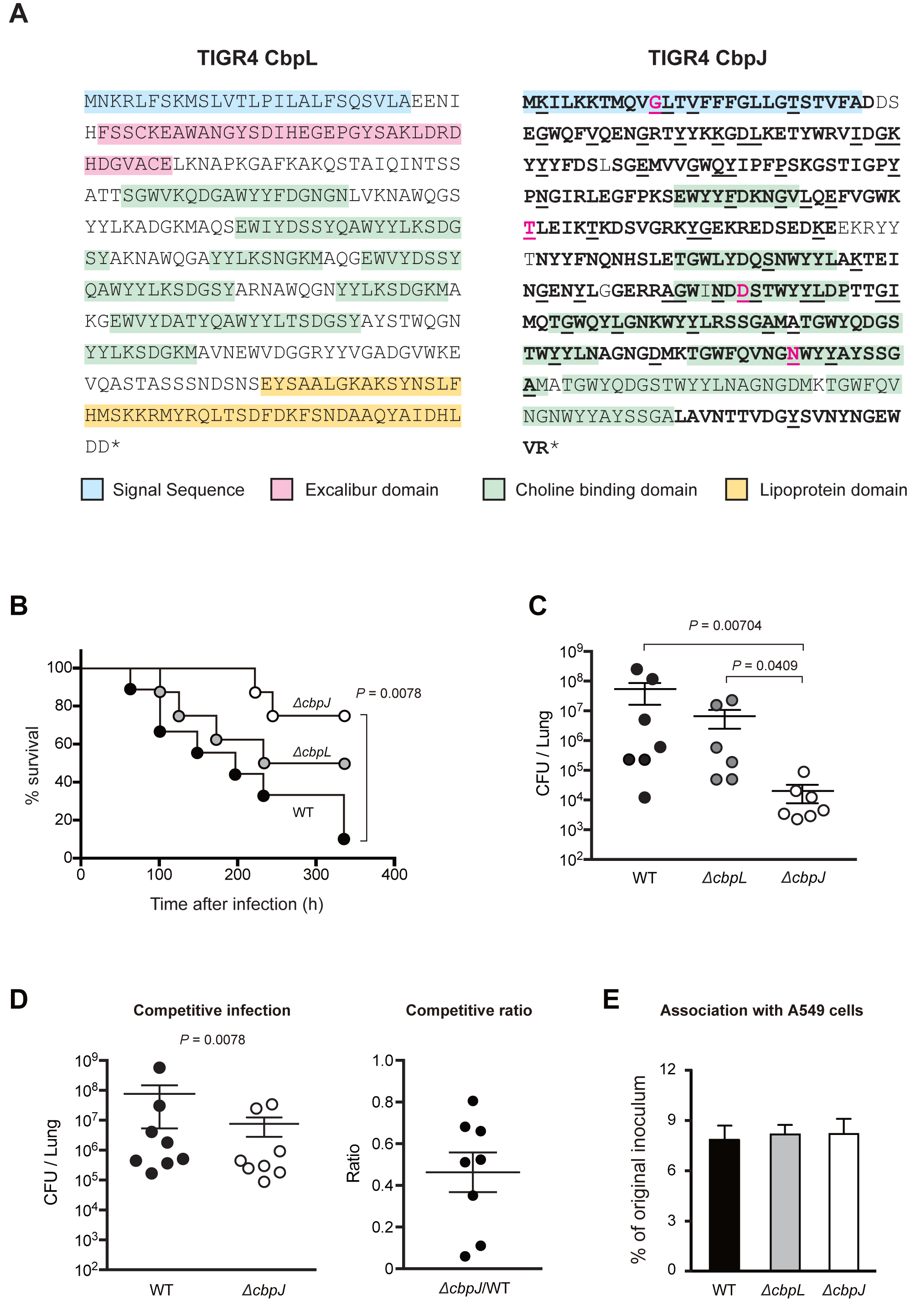
Deficiency of *cbpJ* decreased pneumococcal virulence in mouse pneumonia model. **A.** Amino acid sequences and domain structures of CbpL and CbpJ in strain TIGR4. Bold, black underlined, and magenta underlined characters represent comparable codons and those under purifying or positive selection, respectively. **B.** Mouse pneumonia model. Mice were intranasally infected with 5 × 10^7^ CFU of *S. pneumoniae* TIGR4 WT, *ΔcbpL*, or *ΔcbpJ* strains, and survival was monitored for 14 days. **C.** Pneumococcal CFU in BALF collected at 24 h after intranasal infection. The difference between groups was analysed using the Kruskal-Wallis test with Dunn’ s multiple comparisons test. **D.** *S. pneumoniae* TIGR4 WT and *ΔcbpJ* strains were examined for their competitive infection activities. BALF was collected at 24 h after intranasal infection. The difference between groups was analysed with the Wilcoxon matched-paired signed rank test. **E.** *S. pneumoniae* TIGR4 WT, *ΔcbpL*, and *ΔcbpJ* strains were examined for their ability to associate with A549 cells. Differences between groups were analysed using ordinary one-way ANOVA with Tukey’ s multiple comparisons test. Data are presented as the mean of six samples with standard error (C, D, E).

However, the *S. pneumoniae* WT strain exhibited extensive inflammatory cell infiltration and bleeding compared to that with the *ΔcbpJ* strain. Histological examination of lung tissue from intranasally-infected mice showed that *ΔcbpJ* induced milder inflammation compared to the WT strain. Lung tissue from *ΔcbpL*-infected mice showed moderate inflammation (Fig. 5A). We also measured the bacterial survival rate after incubation with human neutrophils in the absence of serum. Strains *ΔcbpJ* and *ΔcbpL* had a lower survival rate than that of the WT, whereas *ΔcbpJ* showed a slightly increased growth rate compared to that of the WT and *ΔcbpL* strains in RPMI 1640 medium without neutrophils (Fig. 5B and Supplementary Fig. 5D). We also generated recombinant CbpJ using a codon-optimized *cbpJ* sequence for expression in *E. coli* and measured the bacterial survival rate after incubation with neutrophils and the recombinant protein. In the presence of recombinant CbpJ, the survival rate of the *ΔcbpJ* strain was recovered (Supplementary Fig. 6). These results suggest that CbpJ contributes to the evasion of neutrophil-mediated killing. Next, we performed a mouse intravenous infection assay to investigate the role of CbpJ and CbpL in sepsis. In the infection model, the survival rates of *ΔcbpL*- and *ΔcbpJ*-infected mice did not differ significantly from those of mice infected with WT *S. pneumoniae* (Fig. 5C). We also performed a blood bactericidal assay. The survival rates of *ΔcbpJ* and *ΔcbpL* strains in mouse blood were comparable to those of the WT strain (Fig. 5D). We also found that incubation of *S. pneumoniae* in human plasma for 3 h inhibited the expression of *cbpL* and *cbpJ*, as determined by quantitative real-time (q)PCR (Fig. 5E). These results indicate that CbpJ acts as a pneumococcal virulence factor in lung infection by contributing to the evasion of neutrophil-mediated killing, whereas CbpJ has no role in bacterial survival in blood. In addition, *cbpL* deficiency in strain TIGR4 did not significantly attenuate pathogenesis in the mouse lung and blood infection.

**Figure 5:**
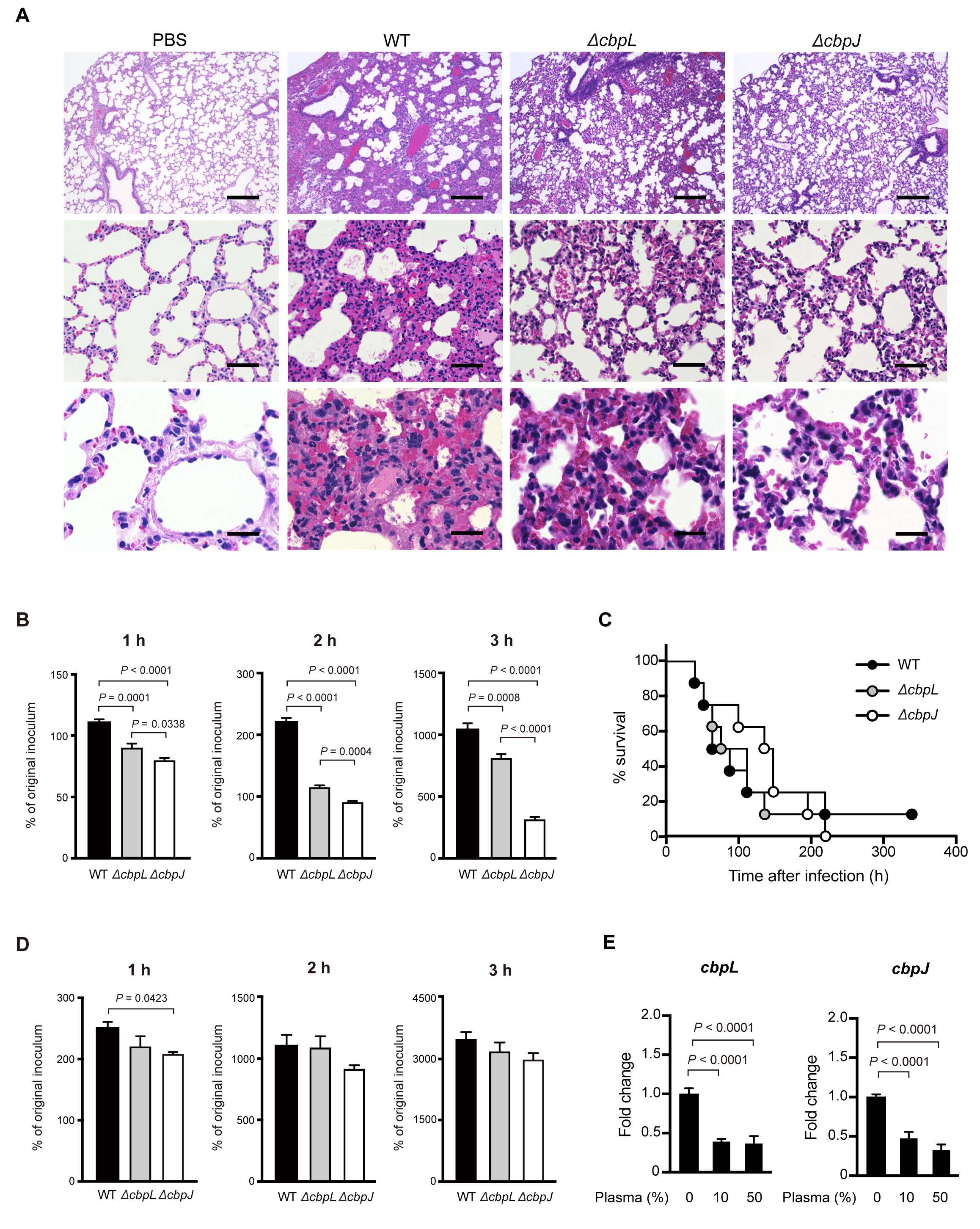
*cbpJ* and *cbpL* are downregulated in the presence of plasma, and do not affect pneumococcal survival in mouse blood. **A.** Haematoxylin and eosin staining of infected mouse lung tissue collected 24 h after intranasal infection with 5 × 10^7^ CFU of *S. pneumoniae* TIGR4 WT, *ΔcbpL*, or *ΔcbpJ* strains. Scale bars, 200 µm (upper panels), 50 µm (middle panels), and 20 µm (lower panels). **B.** Growth of pneumococcal strains in the presence of human neutrophils. Bacterial cells were incubated with neutrophils for 1, 2, and 3 h at 37°C and 5% CO_2_, then serially diluted and plated on TS blood agar. The number of CFUs was determined following incubation. Growth index was calculated by dividing CFU after incubation by the CFU of the original inoculum. Data are presented as the mean of six samples with standard error. **C.** Mouse sepsis model. Mice were intravenously infected with 2 × 10^6^ CFU of *S. pneumoniae* TIGR4 WT, *ΔcbpL*, or *ΔcbpJ*, and survival was monitored for 14 days. Differences between infected mouse groups were analysed with the log-rank test. **D.** Growth of pneumococcal strains in mouse blood. Bacterial cells were incubated in blood for 1, 2, and 3 h at 37°C and 5% CO_2_. Data are presented as the mean of six samples with standard error. **E.** Fold transcript levels of *cbpL* and *cbpJ* in TIGR4 WT *S. pneumoniae* cells in the presence or absence of human plasma. 16S rRNA was used as an internal standard. Data were pooled and normalised from three independent experiments, each performed in quadruplicate.

## Discussion

In this study, we investigated the evolutionarily conserved rates of CBP codons since these cell surface proteins directly interact with the external environment, which induces rapid rates of evolution in genes involved in genetic conflicts^14^. Evolutionary analysis based on phylogenetic relationships can reveal regions in which the encoded amino acids are not allowed to change even under selective pressure. The genetic diversity of *S. pneumoniae* isolated from patients was the result of transmission in a real population. Thus, the evolutionary conservation rate is a parameter that reflects the importance of the protein in human infection. Although so-called arms races involve both the host and bacteria, most studies on genetic diversity have focused on the former^14,27-29^. For example, evolutionary studies based on inter-species comparisons have shown that most of the positive selection targets in host receptors are located in regions that are responsible for direct interactions with pathogens. Our study focused on negative selection targets in bacterial surface proteins through an evolutionary analysis based on intra-species comparisons. This approach enabled us to estimate the contribution of bacterial proteins to species success throughout the life cycle, including inside the host and during the transmission phase.

We previously detected bacterial virulence factors by function prediction – e.g., by searching for conserved motifs/domains, constructing random transposon libraries, or analysing the biochemical properties of the pathogen^30-34^. Although these laboratory-based approaches are valuable, they are time-consuming and costly, and may not yield the expected results. It is useful to examine the correlation between a target molecule and clinical features as this can minimise the time and cost required for analysis. Furthermore, in basic studies on bacterial pathogens, animal infection models are often used to determine whether a bacterial molecule acts as a virulence factor. Although this is the best means of obtaining *in vivo* information, it is unclear how accurately it reflects the clinical condition in humans. Combining an evolutionary analysis and an animal model would thus be highly effective for evaluating the functional significance of a putative virulence factor.

Genome-wide association study (GWAS) is a powerful tool for identifying the relationship between genetic variants – mainly single nucleotide polymorphisms (SNPs) – and phenotype, such as in diseases. As GWAS has become more prevalent, various programs and software packages have been developed for this purpose^35,36^. On the other hand, this approach has certain limitations including the requirement for an appropriate control group and detailed information regarding phenotype. In infectious diseases, it can be difficult to quantify clinical features recorded at different medical centres. Furthermore, in the case of most pathogens, there are no natural attenuated or avirulent strains that can serve as a control group. Our evolutionary analysis has the advantage that it can be performed with genomic information of pathogenic strains only by assuming the presence of pathogens as a phenotype evading natural selection. Since synonymous and non-synonymous substitutions are estimated to occur with equal probability under no selective pressure, a population in which the latter has resulted in extinction by natural selection can serve as a control group. While we have shown in the current study that evolutionary analysis with a small population has the power to detect evolutionarily conserved proteins, a larger population would allow a higher-resolution analysis, including detection of conserved regions in some pathogenic strains isolated from a specific site of infection or pathological condition. Since this analysis involves simultaneous processing of aligned nucleotide and amino acid sequences, more information is obtained from only SNPs extracted from nucleotide sequences. In addition, automated phylogenetic and evolutionary analyses are needed to analyse a large population. Therefore, the development of software packages for meta-data is expected to aid the widespread application of this analytical approach.

There are some limitations to our evolutionary analysis. Firstly, although it can detect evolutionarily conserved proteins, it cannot identify diverse virulence factors such as PspA and CbpA within species^19,37,38^. Similarly, virulence factors recently acquired by horizontal gene transfer have not been under selective pressure for a sufficiently long period to perform this analysis. In addition, the high rate of codons under negative selection indicate their universal importance in bacterial species. In other words, a molecule under relaxed selective pressure could contribute to the virulence of some populations of the species. However, these features of molecular evolutionary analysis can be advantages when screening for therapeutic target sites or vaccine antigens with a low frequency of missense mutations, which could reduce the virulence or survivability of the pathogen. Evolutionary analysis could also be an effective alternative strategy for overcoming drug resistance through antigen replacement, and could reduce costs associated with drug discovery and development.

The *lytA* gene, which was conserved among virtually all pneumococcal strains, showed the highest rates of codons under negative selection, except for *cbpJ* that was only present in some strains. LytA is known to induce pneumococcal-specific autolysis^39^ and contributes to pneumococcal virulence^16,40^. Our evolutionary analysis supports previous reports that *lytA* is a suitable genetic marker^41,42^ due to its evolutionary conservation. We also showed that *pspA* and *cbpA* show relatively high rates of codons under positive selection, and both encode polymorphic virulent proteins^17,19,37^ that are candidate vaccine antigens, even though these genes are not universally present within a global serotype 1 collection^38^. In addition, selective pressure by vaccines can easily cause differentiation or deficiency of these proteins as the corresponding genes contain few codons under negative selection. A multivalent system would be required for vaccines prepared using these antigens.

An *in vivo* competition assay in mice indicated that deficiency of *cbpJ* is a disadvantage for pneumococcal survival *in vivo.* On the other hand, co-infection showed a smaller difference in bacterial CFUs between WT and *ΔcbpJ* as compared to each single infection. In the single infection of the *ΔcbpJ* strain, the bacteria could not be protected by CbpJ. However, in co-infection, the interaction of neutrophils and CbpJ in the WT strain could suppress neutrophil killing activity. In addition, some CbpJ may be released from the WT strain by autolysis. As a result, some of the *ΔcbpJ* strain could have been protected similar to the WT strain. Concerning selection, it was previously reported that a single cell bottleneck effect in pneumococcal infection occurs during bloodstream invasion and in transmission between hosts^43,44^. Our finding also suggests that a bottleneck effect occurs in a limited situation. The difference in bacterial burden of BALF between single and competitive infections suggested a possibility that the bottleneck effect plays a more important role for the selection of *cbpJ*-lacking cells compared to the competition in the lung.

In this study, *cbpL* and *cbpJ* were downregulated in the presence of plasma. Although regulation of CBPs is still largely unknown, one possible hypothesis is that the genes are regulated by a pneumococcal two component system (TCS). *S. pneumoniae* interplays with its environment by using 13 TCSs and one orphan response regulator^45,46^. TCSs typically consist of a membrane-associated sensory protein called a histidine kinase and a cognate cytosolic DNA-binding response regulator, which acts as a transcriptional regulator. Although specific stimuli to histidine kinases still remain unclear, there is a possibility that a histidine kinase sensor protein of the TCSs can respond to some plasma components.

Although the difference was not statistically significant, mice intranasally infected with TIGR4 *ΔcbpL* strain showed a trend towards improved survival relative to the WT-infected mice. In a previous study, a D39*lux cbpL*-deficient strain showed reduced virulence compared to the WT strain^22^. Since CbpL sequences in TIGR4 and D39 strains are similar, the discrepancy between the previous study and our findings is likely due to differences in other surface proteins in each strain. For example, the absence of CbpJ, which contributes to the evasion of neutrophil killing, could affect the survivability of D39.

Frolet *et al.* reported that both CbpJ and CbpL are considered as possible adhesins because they display interaction with C-reactive protein (CRP), and CRP, elastin, and collagen in solid phase assay, respectively^47^. Meanwhile, Gosink *et al.* showed no significant differences in Detroit nasopharyngeal cells adhesion, rat nasopharynx colonization, and pathogenesis in the sepsis model between the WT and the *cbpJ* mutant strains^48^. Their results are mostly consistent with our data. We also showed that there were no significant differences in the A549 cells adhesion assay and in intravenous infection as a sepsis model. On the other hand, we found a difference in the lethal intranasal mouse infection that is completely different from the non-lethal colonization model. We consider that CbpJ contributes to pneumococcal evasion of host immunity rather than colonization. Concerning CbpL, elastin and collagen are extracellular matrix proteins and binding activity to these proteins could contribute to bacterial adhesion, whereas CRP is found in blood plasma and is used as a marker of inflammation. However, CbpL did not contribute at least to pneumococcal adhesion to A549 cells. There is a discrepancy between protein-protein interactions in the solid phase and host cell-bacteria interactions.

Recently, anti-virulence drugs have been developed as an additional strategy to treat or prevent bacterial infections. Drugs targeting bacterial virulence factors are expected to reduce the selective pressure of conventional antibiotics since they would not affect the natural survival of targeted bacteria^49^. Furthermore, the abundance of candidate targets is a major advantage of antivirulence strategies. Effective design of vaccines and antivirulence drugs requires a thorough understanding of virulence factors; combining our evolutionary analysis and traditional molecular microbiological approaches can improve the detection of potential drug targets. In this study, we identified CbpJ as a novel evolutionarily conserved virulence factor. Thus, molecular evolutionary analysis is a powerful system that can reveal the importance of virulence factors in real-world infections and transmission.

## Methods

### Phylogenetic and evolutionary analyses

Phylogenetic and evolutionary analyses were performed as described previously ^50,51^, with minor modifications. Homologues and orthologues of *cbp* genes were searched using the tBLASTn function of NCBI BLAST. Domain structures of CbpJ and CbpL were searched by MOTIF Search^23^ with PROSITE, NCBI-CDD, and P-fam^24-26^. Bacterial ORFs and promoters were predicted by FGENESB (Bacterial Operon and Gene Prediction) and BPROM, respectively^52^. To prevent node density artefacts, sequences with 100% identity were treated as the same sequence in Phylogears2^53,54^. The sequences were aligned using MAFFT v.7.221 with an L-INS-i strategy^55^, and ambiguously aligned regions were removed using Jalview^56,57^. Calculated orthologous regions were used for further phylogenetic analysis, and edited codon sequences were re-aligned using MAFFT with an L-INS-i strategy. The best-fitting codon evolutionary models for MrBayes and RAxML analyses were determined using Kakusan4^58^. Bayesian Markov chain Monte Carlo analyses were performed with MrBayes v.3.2.5^59^, and 2 × 10^6^ generations were sampled after confirming that the standard deviation of split frequencies was < 0.01 for up to 8 × 10^6^ generations. To validate phylogenetic inferences, maximum likelihood phylogenetic trees with bootstrap values were generated with RaxML v.8.1.20^60^. Phylogenetic trees were generated using FigTree v.1.4.2^61^ based on the calculated data.

Evolutionary analyses were performed based on aligned orthologous regions of *cbp* genes and Bayesian phylogenetic trees. Whole-gene non-synonymous/synonymous ratio calculations as well as statistical tests for negative or positive selection of individual codons were performed using the two-rate fixed-effects likelihood function in HyPhy software package^62^.

### Bacterial strains and construction of mutant strains

*Streptococcus pneumoniae* strains were cultured in Todd-Hewitt broth (BD Biosciences, Franklin Lakes, NJ, USA) supplemented with 0.2% yeast extract (BD Biosciences) (THY medium) at 37°C. For mutant selection and maintenance, spectinomycin (Wako Pure Chemical Industries, Osaka, Japan) was added to the medium at a concentration of 120 µg/ml.

*S. pneumoniae* TIGR4 isogenic *cbpJ* (*ΔcbpJ*) and *cbpL* (*ΔcbpL*) mutant strains were generated as previously described^33^. Briefly, the upstream region of *cbpJ* or *cbpL*, an *aad9* cassette, and the downstream region of *cbpJ* or *cbpL* were combined by PCR using the primers shown in Supplementary Table 4. The products were used to construct the mutant strains by double-crossover recombination with the synthesised CSP2^63^. All mutations were confirmed by PCR amplification of genomic DNA isolated from the mutant strains. For growth measurements, pneumococci were cultured until the optical density at 600 nm (OD_600_) reached 0.4, and the exponential phase cultures of each strain were back-diluted into fresh THY and grown at 37°C. Growth was monitored by measuring the values of OD_600_ every 0.5-1 hour. For the following assays, *S. pneumoniae* strains were grown to exponential growth phase (OD_600_ = ∼0.4) unless otherwise indicated, and then resuspended in PBS or the appropriate buffer.

### Preparation of recombinant CbpJ

The *cbpJ* sequence without codons encoding the signal peptide sequence was optimized for *E. coli* using GENEius software, and the optimized sequence was synthesized (Eurofins Genomics, Brussel, Belgium). Optimized *cbpJ* and pQE-30 vector (Qiagen, Valencia, CA, USA) were amplified with the specific primers listed in Supplementary Table 5 and PrimeSTAR^®^ MAX DNA Polymerase (TaKaRa Bio, Shiga, Japan). The DNA fragments were assembled using the GeneArt^®^ Seamless Cloning and Assembly Kit (Thermo Fisher Scientific, Waltham, MA, USA). The constructed plasmid was transformed into *E. coli* XL-10 Gold (Agilent, Santa Clara, CA, USA), and recombinant CbpJ was purified as described previously^31,33,64-66^.

### Blood and neutrophil bactericidal assays

A blood bactericidal assay was performed as previously described^31,33,67^. Mouse blood was obtained via cardiac puncture from healthy female CD-1 mice (Slc:ICR, 6 weeks old; Japan SLC, Hamamatsu, Japan). For human neutrophil isolation, blood was collected via venepuncture from healthy donors after obtaining written, informed consent according to a protocol approved by the institutional review board of Osaka University Graduate School of Dentistry (H26-E43). Neutrophils were isolated from fresh human blood by density gradient centrifugation using Polymorphprep (Alere Technologies, Jena, Germany). Pneumococcal cells grown to the mid-log phase were washed and resuspended in phosphate-buffered saline (PBS). Bacterial cells (1 × 10^4^ CFU/20 µl) were combined with fresh mouse blood (180 µl) or human neutrophils (2 × 10^5^ cells/180 µl) in RPMI 1640 medium, and the mixture was incubated at 37°C with 5% CO_2_ for 1, 2, and 3 h. Viable cell counts were determined by seeding diluted samples onto THY blood agar. The percent of the original inoculum was calculated as the number of CFU at the specified time point divided by the number of CFU in the initial inoculum.

### MIC and MBC assays

Minimum inhibitory concentration (MIC) and minimum bactericidal concentration (MBC) assays were performed as previously described^51,68^. For MIC and MBC assays, 0.5-1.0 × 10^4^ bacteria were added into THY broth supplemented with 2-fold serial dilutions of penicillin G. Bacterial growth after 24 hours at 37°C in anaerobic conditions was spectrophotometrically measured at OD_600_. We defined the OD_600_ values less than 0.06 as complete inhibition of bacterial growth. To determine MBCs, we inoculated 5 µL of the bacterial cultures onto TS blood agar and incubate them at 37°C in anaerobic conditions. The antimicrobial concentration at which no growth was detectable was defined as the MBC.

### Mouse infection assays

All mouse experiments were conducted in accordance with animal protocols approved by the Animal Care and Use Committee of Osaka University Graduate School of Dentistry (28-002-0). Female CD-1 mice (Slc:ICR, 6 weeks old) were intranasally infected with 5 × 10^7^ or 2 × 10^6^ CFU of *S. pneumoniae* via the tail vein. Mouse survival was monitored for 14 days. At 24 h after intranasal infection, animals were euthanized by lethal intraperitoneal injection of sodium pentobarbital and lung tissue or BALF samples were collected. Bacterial counts in BALF were determined by plating serial dilutions. Lung tissue specimens were fixed with 4% formaldehyde, embedded in paraffin, and cut into sections that were stained with haematoxylin and eosin solution (Applied Medical Research, Osaka, Japan) and visualized with a BZ-X710 microscope (Keyence, Osaka, Japan). For the competition assay, CD-1 mice were intranasally infected with 20 µL of the mixture of wild-type (1.0 × 10^7^ CFU) and *ΔcbpJ* (1.5 × 10^7^ CFU) strains resuspended in PBS, in total, ∼2.5 × 10^7^ CFU. BALF samples were collected at 24 h after infection and bacterial counts in BALF were determined. Total and mutant strain CFUs were determined by serial dilution plating on TS blood agar with or without spectinomycin. The CFU number for the wild-type strain was calculated by subtracting that of the mutant strain from the total CFUs.

### qPCR

qPCR was performed as previously described^50,51^, with minor modifications. Primers are listed in Supplementary Table 4. Total RNA of pneumococcal strains grown to the mid-log phase (OD_600_ = 0.4-0.5) was isolated with an RNeasy Mini kit (Qiagen) and RQ1 RNase-Free DNase (Promega, Madison, WI, USA), and cDNA was synthesised with SuperScript IV VILO Master Mix (Life Technologies, Carlsbad, CA, USA). qPCR analysis was performed on a StepOnePlus Real-Time PCR system using Power SYBR Green Master PCR mix (Thermo Fisher Scientific). 16S rRNA was used as a normalising control.

### Statistical analysis

Statistical analysis of *in vitro* and *in vivo* data was performed with Mann-Whitney test, Kruskal-Wallis test with Dunn’s multiple comparisons test, Wilcoxon matched-paired signed rank test, and ordinary one-way ANOVA with Tukey’s multiple comparisons test. Mouse survival curves were compared with the log-rank test. *P* < 0.05 was considered statistically significant. The tests were performed on Prism v.6.0h or v.7.0d software (GraphPad Inc., La Jolla, CA, USA). All experiments were repeated at least three times. In the evolutionary analyses, *P* < 0.1 was regarded as a significant difference with the HyPhy default setting.

## Supporting information

## Acknowledgements

This study was supported in part by the Japan Society for the Promotion of Science KAKENHI (grant numbers 15H05012, 16H05847, 16K15787, 17H05103, 17K11666, and 17H04369); SECOM Science and Technology Foundation; Takeda Science Foundation; GSK Japan Research Grant; Asahi Glass Foundation; Kurata Memorial Hitachi Science and Technology Foundation; and Kobayashi International Scholarship Foundation. The funders had no role in study design, data collection or analysis, decision to publish, or preparation of the manuscript.

## Author contributions

M.Y. and S.K. designed the study. M.Y. and Y.Y. performed bioinformatics analyses. K.G., M.Y., and Y.H. performed the experiments. M.Y., T.S., M.N., and S.K. contributed to the setup of the experimentation. M.Y. wrote the manuscript. G.K., Y.H., Y.Y., T.S., M.N., K.N., and S.K. contributed to the writing of the manuscript.

