## Supplementary material for "Identification of evolutionarily conserved virulence factor by selective pressure analysis of *Streptococcus pneumoniae*"

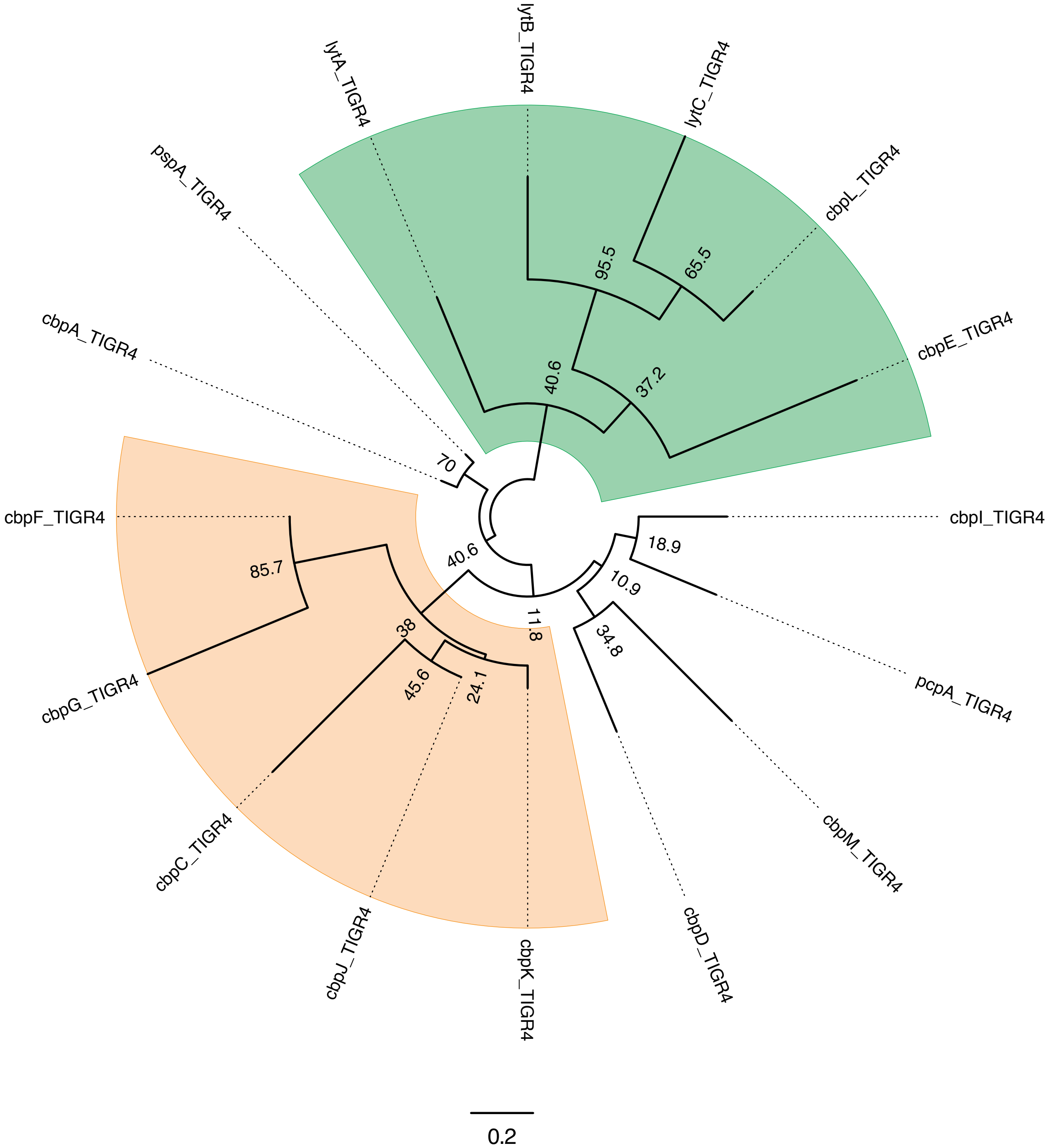


**Supplementary Figure 1. Maximum likelihood phylogenetic analyses of *cbp* genes in *S. pneumoniae* strain TIGR4.** Nucleotide-based maximum likelihood phylogenetic tree of *cbp* genes in TIGR4. The tree is unrooted and bootstrap values are shown near the nodes. The scale bar indicates nucleotide substitutions per site.

Figure S1. Yamaguchi *et al.*

**A**

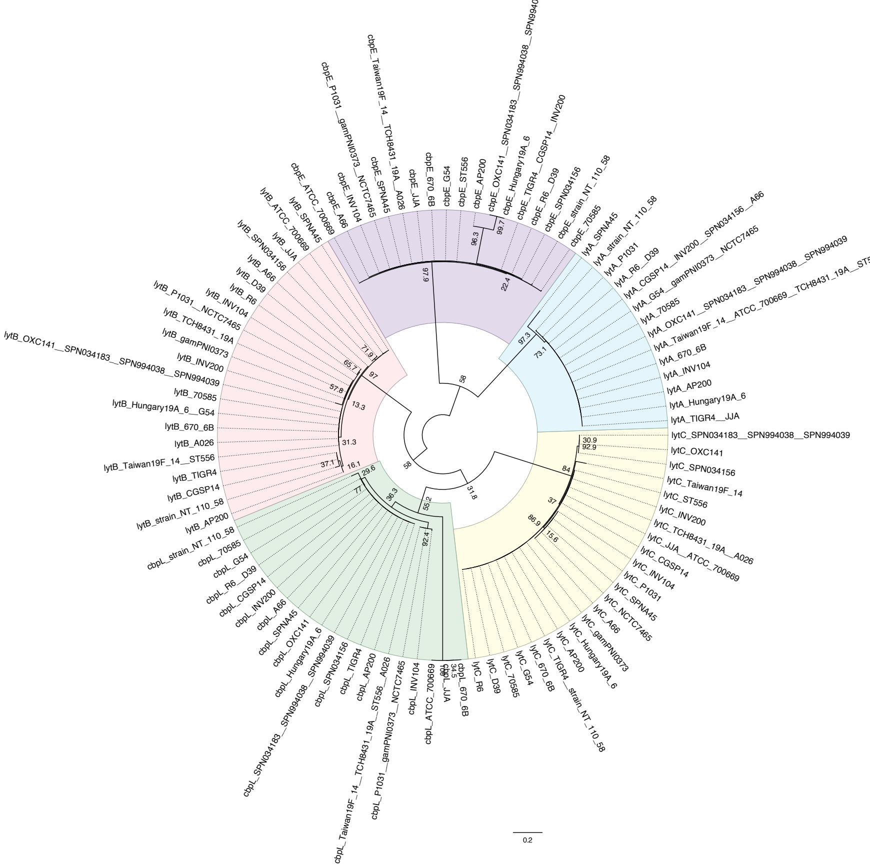

**B**

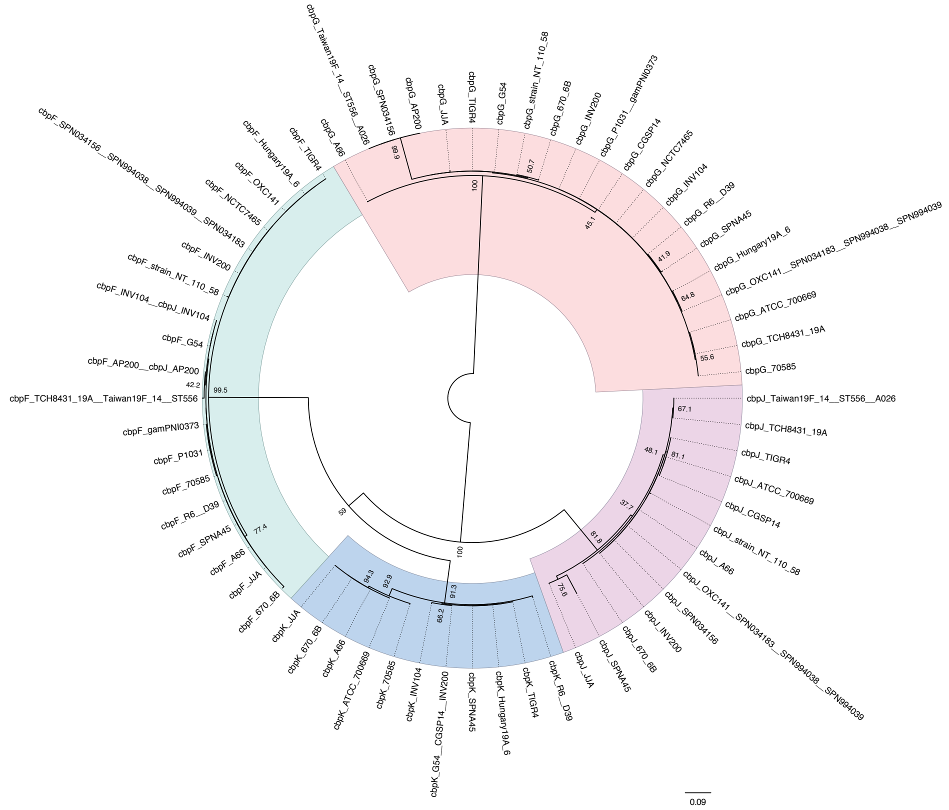

**Supplementary Figure 2. Maximum likelihood phylogenetic analyses of *cbp* genes with high similarity. A, B.** Nucleotide-based maximum likelihood phylogenetic tree of the *lytA*, *lytB*, *lytC*, *cbpE*, and *cbpL* genes (A) and *cbpF*, *cbpG*, *cbpJ*, and *cbpK* genes (B) in *S. pneumoniae*. The trees are unrooted but are presented as midpoint-rooted for clarity. Strains with identical sequences are shown on the same branch, and bootstrap values are shown near the nodes. The scale bar indicates nucleotide substitutions per site.

Figure S2. Yamaguchi *et al.*

A

B

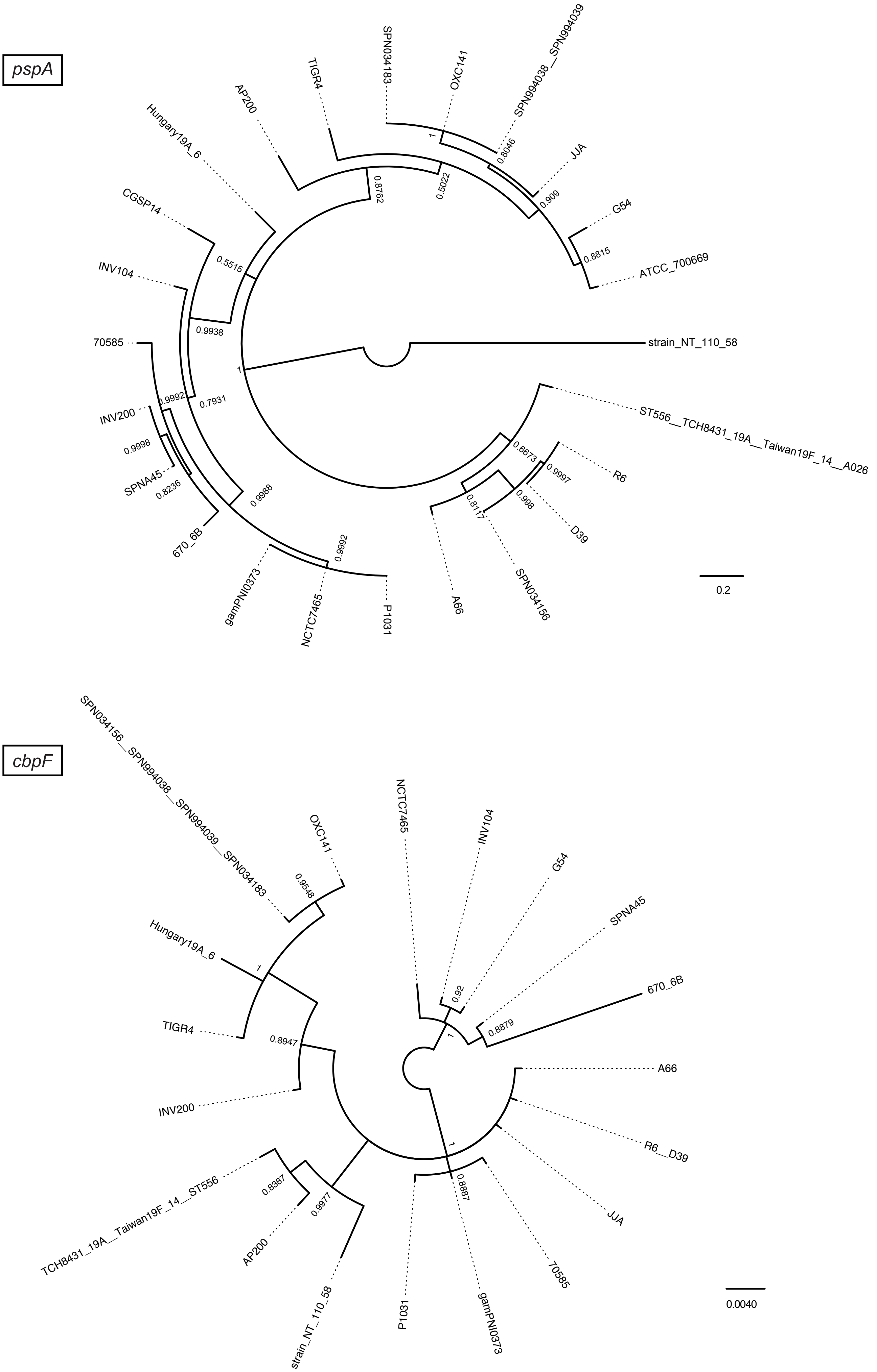


Figure S3. Yamaguchi *et al.*

C

*cbpJ*

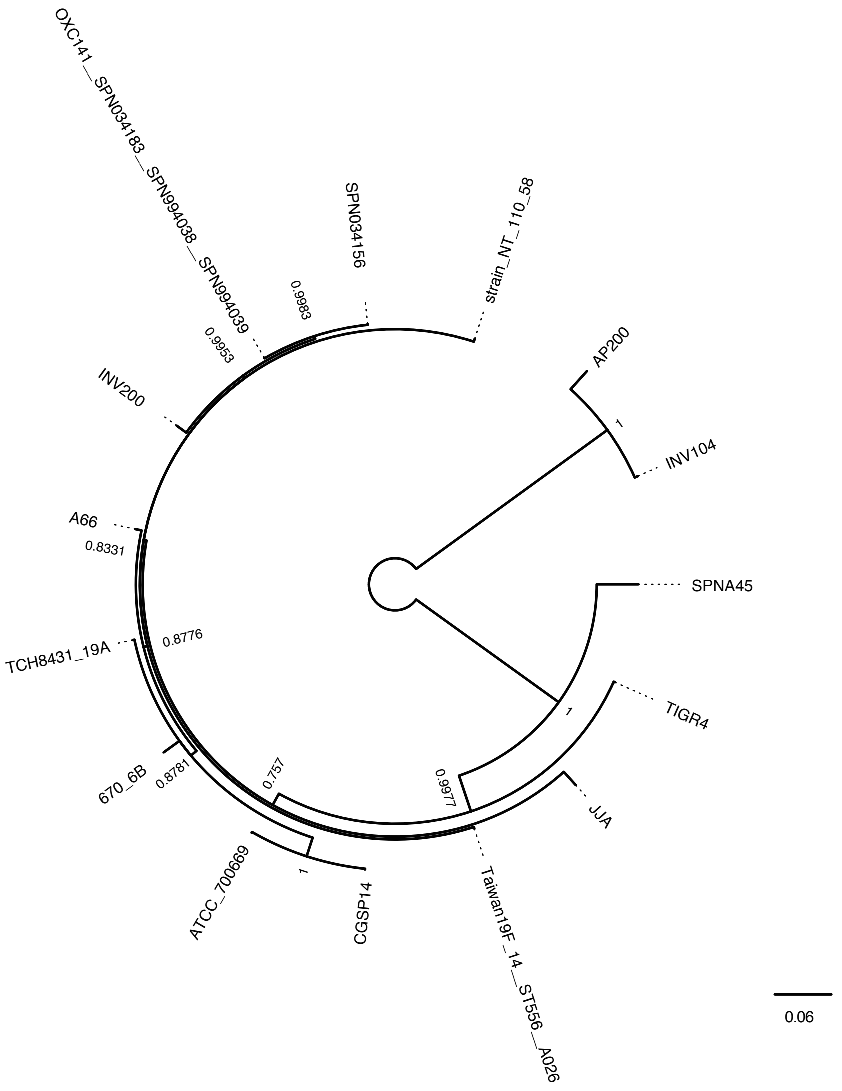


D

*cbpK*

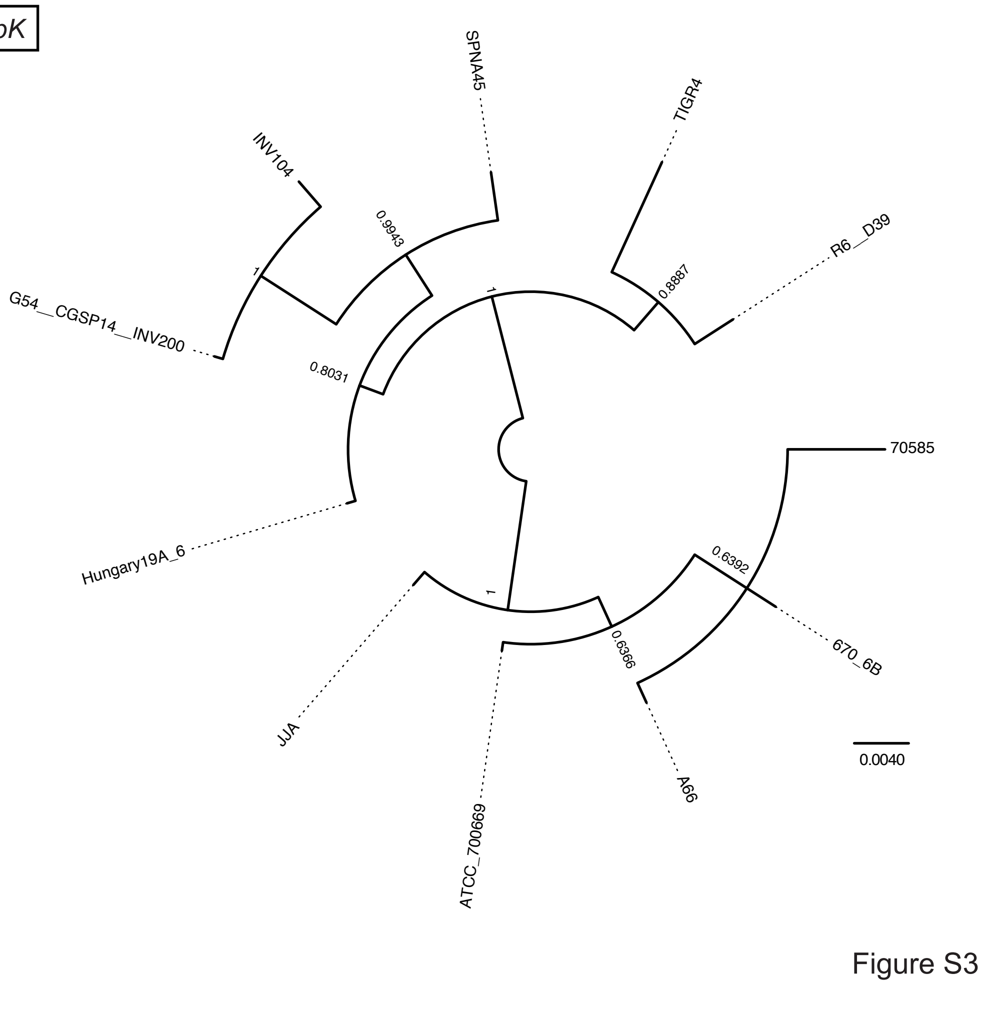


Figure S3. Yamaguchi *et al.*

E

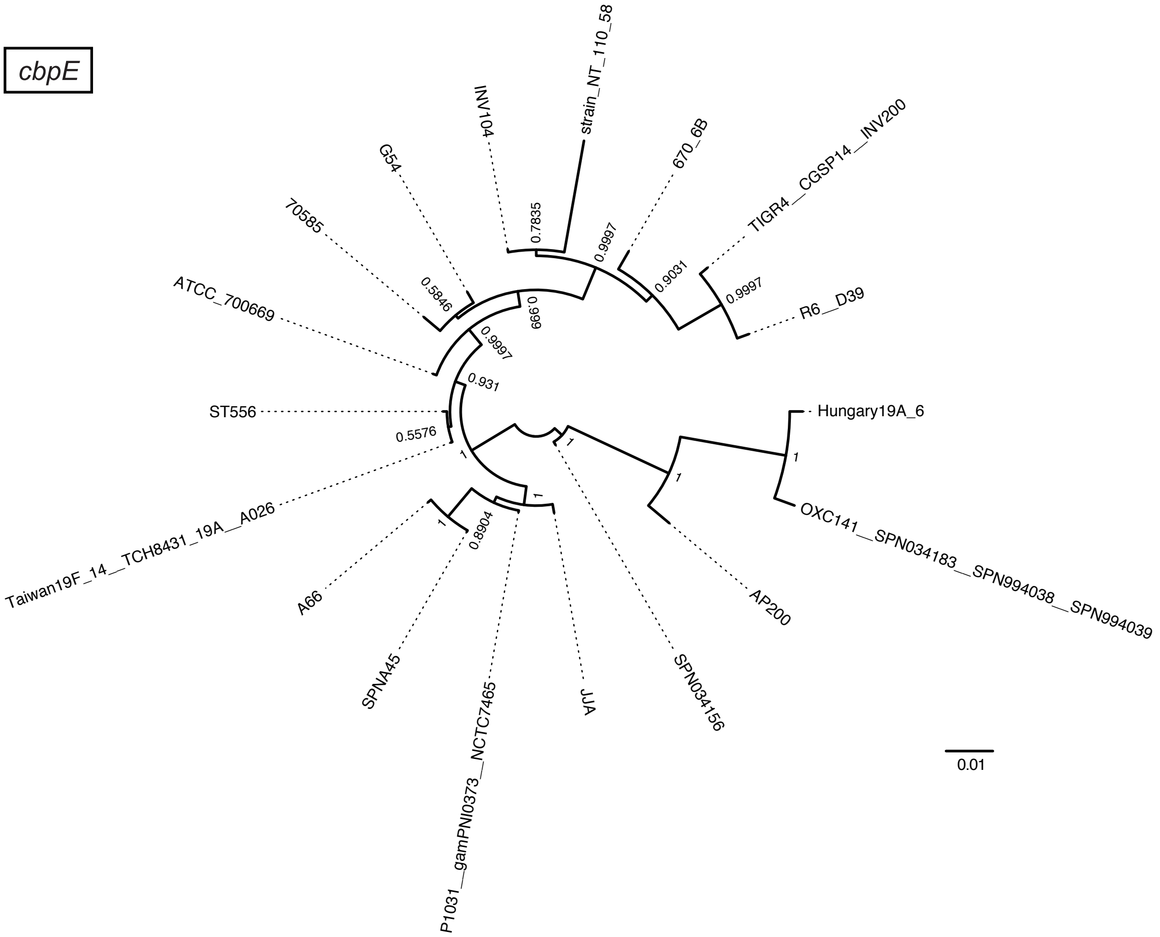


F

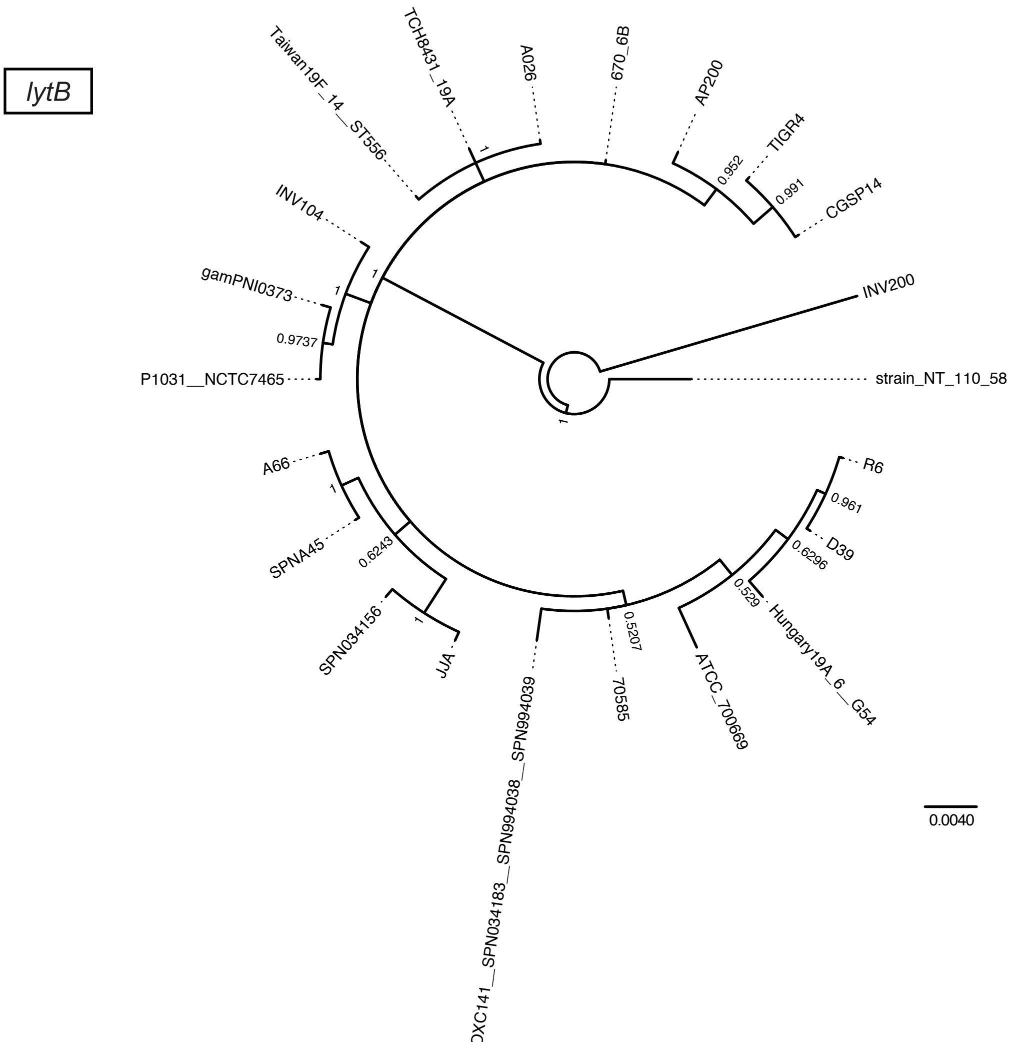


Figure S3. Yamaguchi *et al.*

G

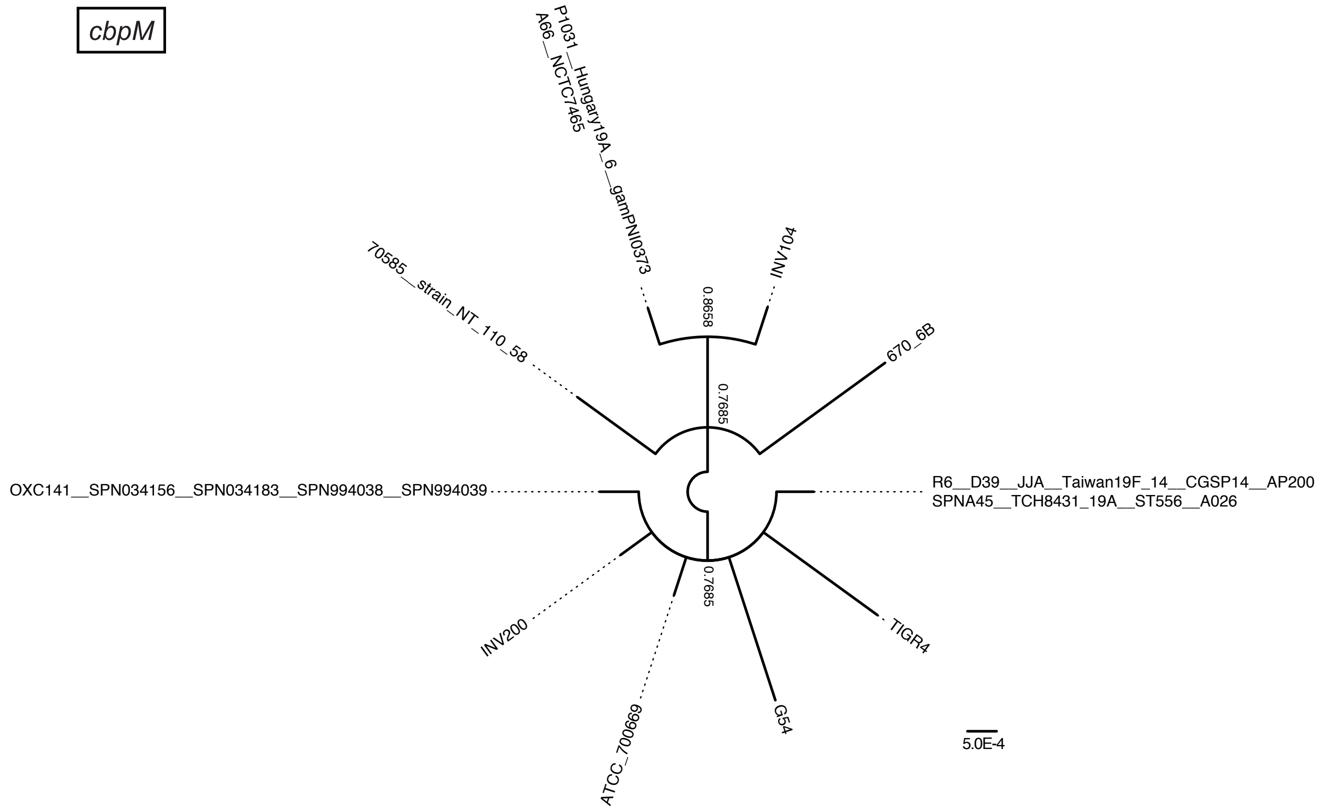


H

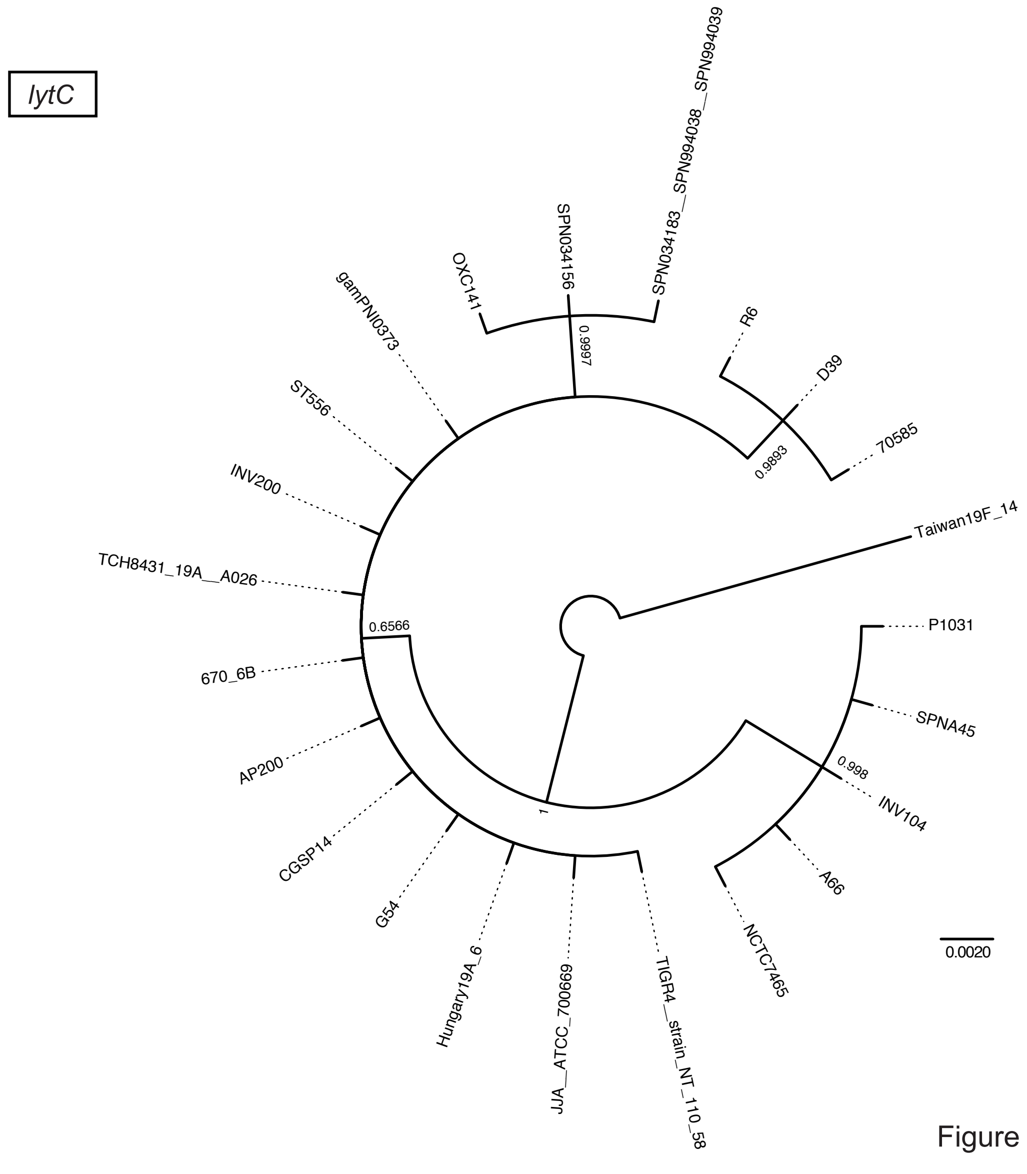


Figure S3. Yamaguchi *et al.*

I

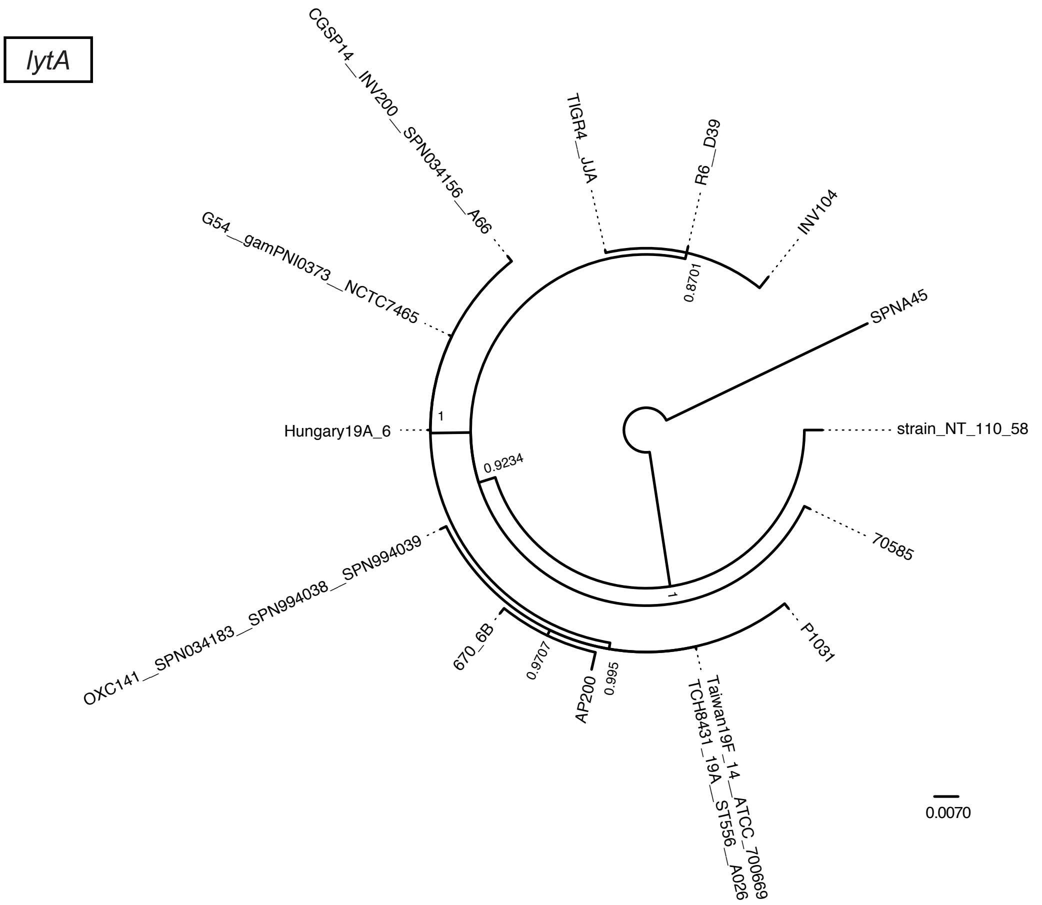


J

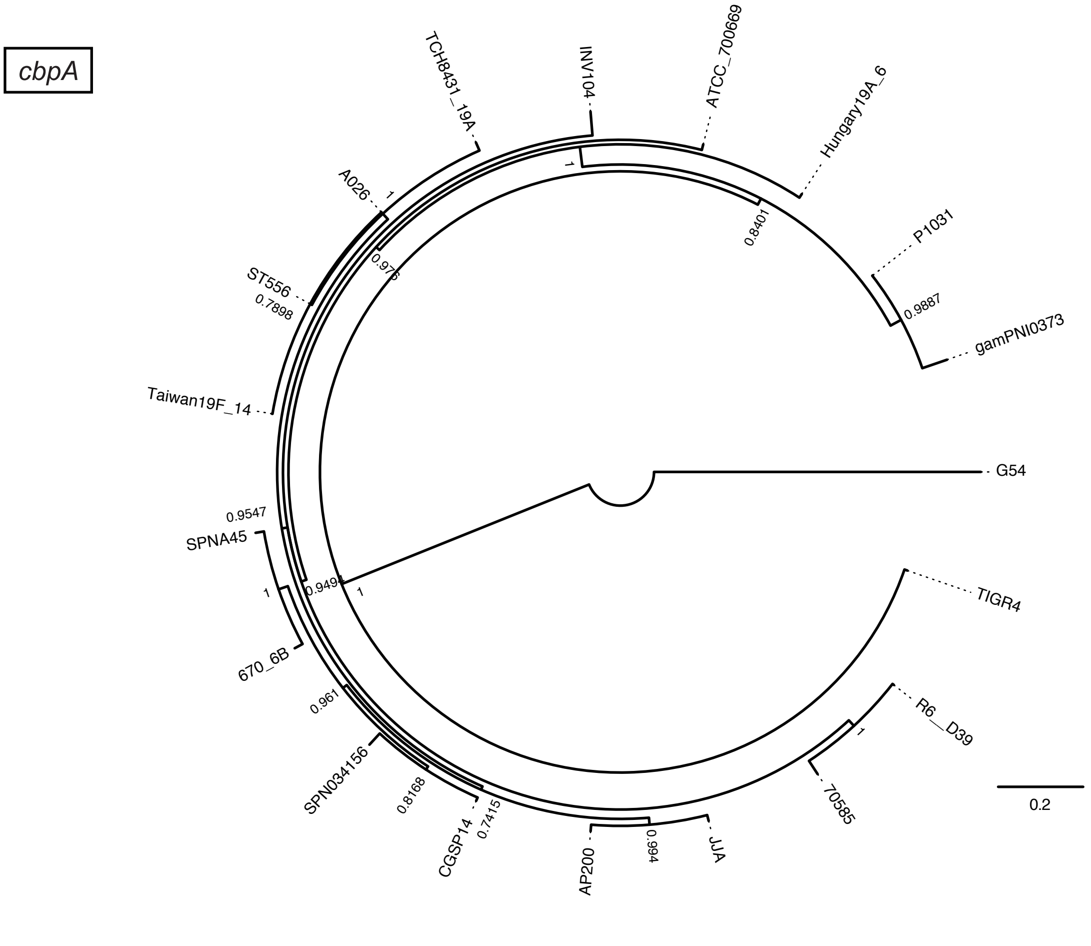


Figure S3. Yamaguchi *et al.*

**Supplementary Figure 3. Codon-based Bayesian phylogenetic tree of genes encoding CBPs in *S. pneumoniae*.** Posterior probabilities are shown near the nodes. Strains with identical sequences are shown on the same branch. The scale bar indicates nucleotide substitutions per site. The trees are unrooted but are presented as a midpoint-rooted polar tree for clarity. Each figure shows the phylogenetic tree of the following genes: A, *pspA*; B, *cbpF*; C, *cbpJ*; D, *cbpK*; E, *cbpE*; F, *lytB*; G, *cbpM*; H, *lytC*; I, *lytA*; and J, *cbpA*.

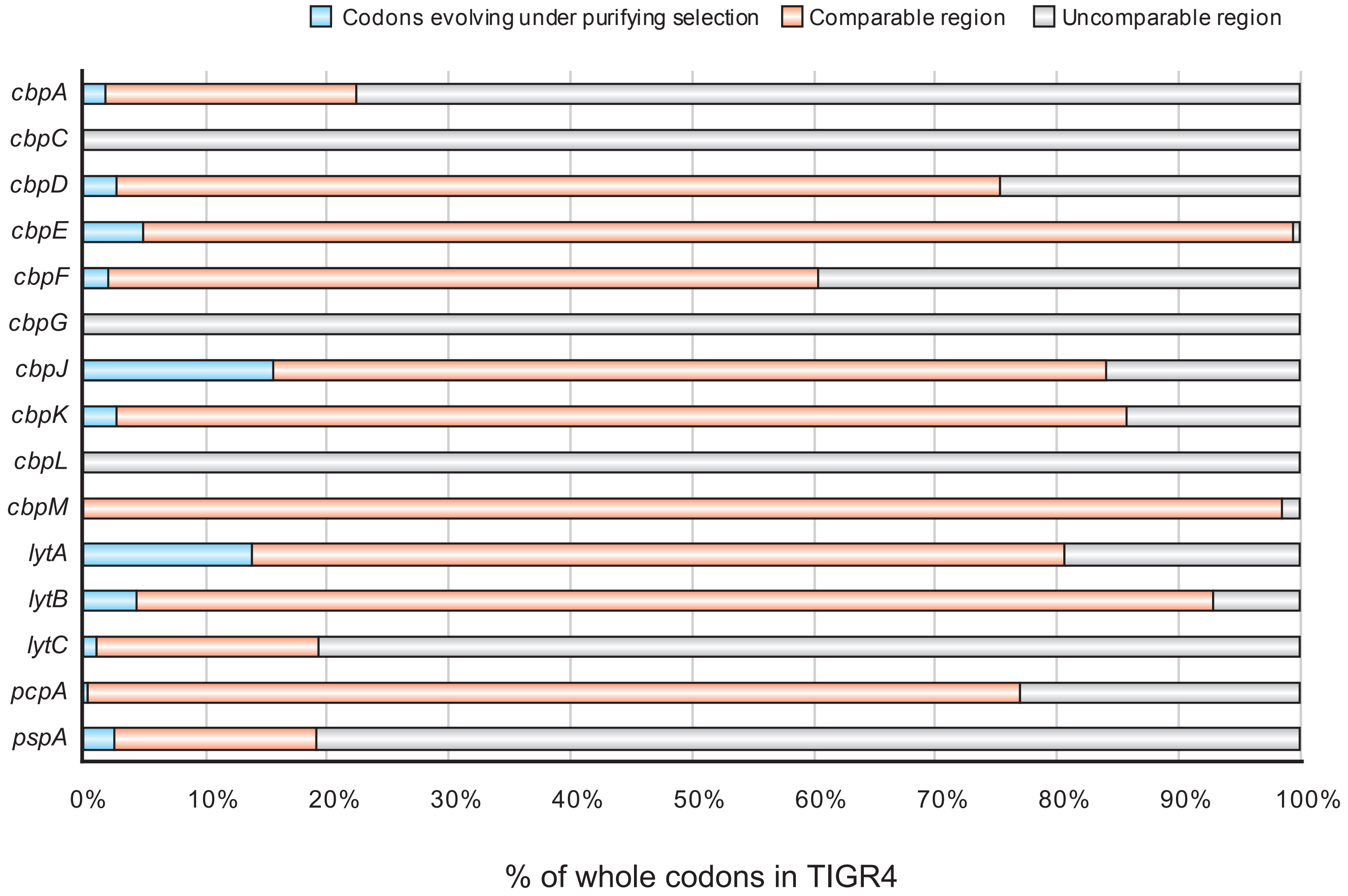


**Supplementary Figure 4. Rate of evolutionarily conserved codons in pneumococcal CBPs.** Codons selection of pneumococcal CBPs under purifying was identified with HyPhy software using phylogenetic trees and aligned sequences. Blue, orange, and grey represent the rate of codons under purifying selection, comparable common codons, and incomparable codons, respectively. Actual numbers and other parameters are shown in Table 1.

Figure S4. Yamaguchi *et al.*

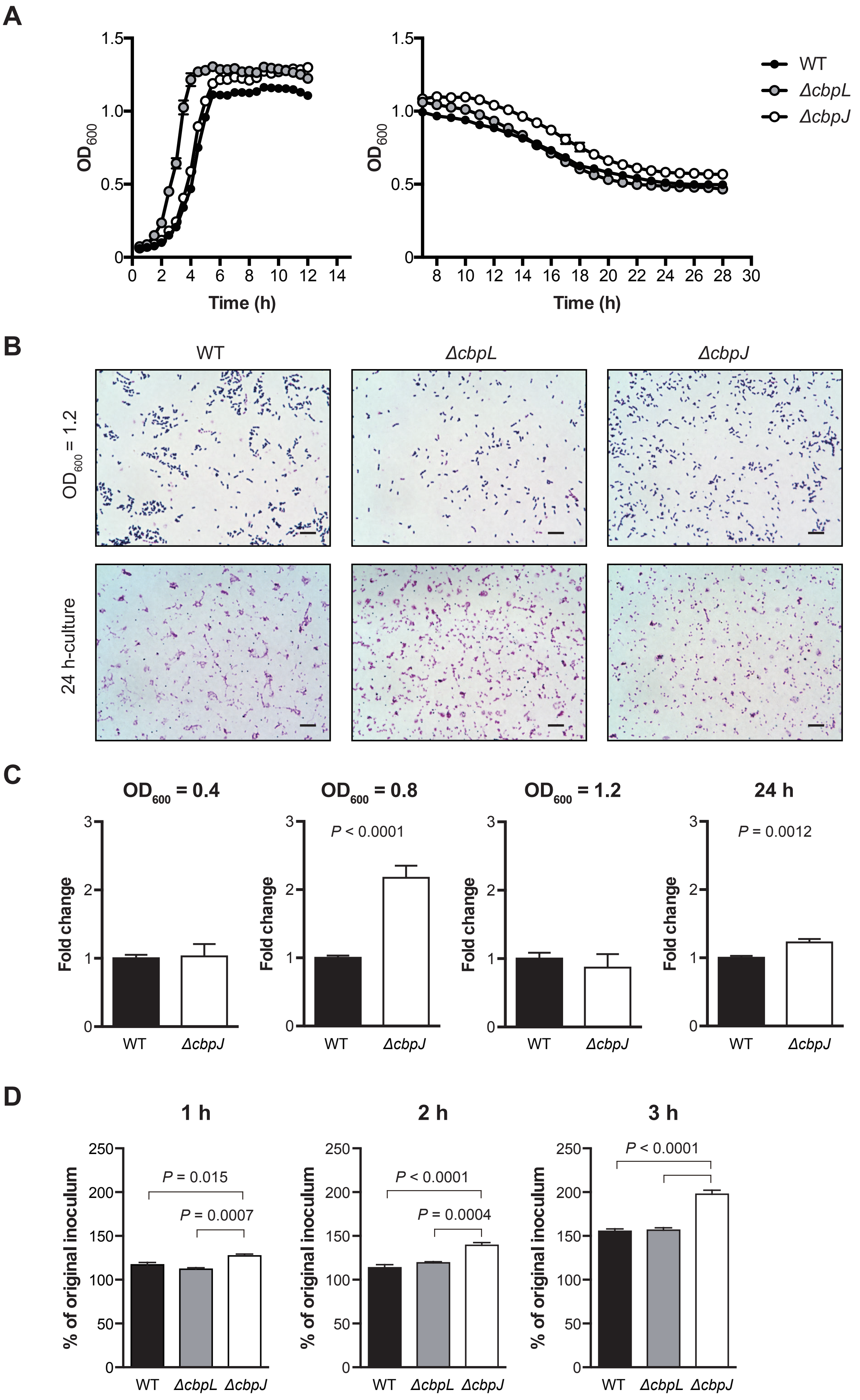


Figure S5. Yamaguchi *et al.*

**Supplementary Figure 5. A.** ***S. pneumoniae*** **strains were grown in THY broth at 37˚C.** Data are presented as the means of 5 samples from a representative experiment. S.E. values are represented by vertical lines. **B.** Gram staining of *S. pneumoniae* TIGR4 WT, *ΔcbpL*, or *ΔcbpJ* strains in THY broth. The scale bars indicate 10 µm. **C.** Fold transcript levels of *lytA* in *S. pneumoniae* TIGR4 WT and *ΔcbpJ* strains in THY broth. 16S rRNA was used as an internal standard. Data were pooled and normalised from three or four independent experiments, each performed in triplicate. **D.** Growth of pneumococcal strains in RPMI 1640 medium. Bacterial cells were incubated for 1, 2, and 3 h at 37°C and 5% $CO_2$, then serially diluted and plated on TS blood agar. The number of CFUs was determined following incubation. Differences between groups were analysed using ordinary one-way ANOVA with Tukey's multiple comparisons test.

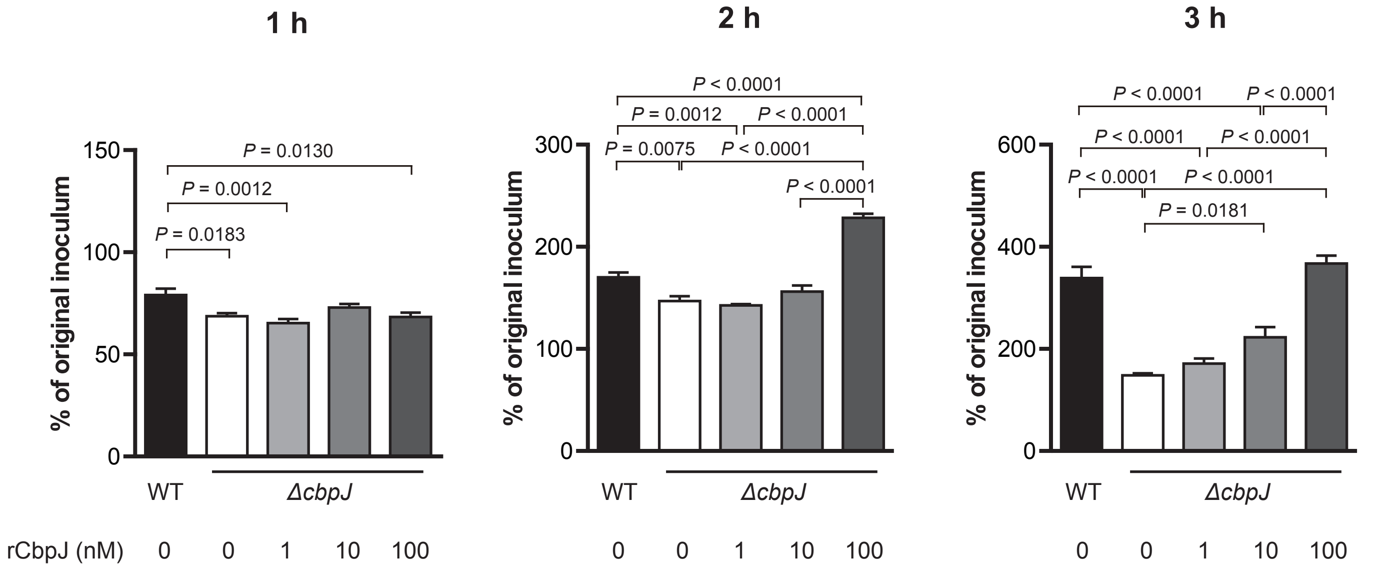


**Supplementary Figure 6. Growth of pneumococcal strains with rCbpJ in the presence of human neutrophils.** Bacterial cells were incubated with rCbpJ and neutrophils for 1, 2, and 3 h at 37°C and 5% $CO_2$, then serially diluted and plated on TS blood agar. The number of CFUs was determined following incubation. Data are presented as the mean of six samples with standard error. Differences between groups were analysed using ordinary one-way ANOVA with Tukey's multiple comparisons test.

**Supplementary Table 1.** Locus tag number of *cbp* genes in each pneumococcal strain

| | TIGR4 | R6 | D39 | 70585 | JJA | P1031 | Taiwan19F-14 | Hungary19A-6 | G54 | CGSP14 | ATCC 700669 | AP200 | 670-6B | SPNA45 |
|---|---|---|---|---|---|---|---|---|---|---|---|---|---|---|
| *cbpA* | SP_2190 | spr1995 | SPD_2017 | SP70585_2316 | SPJ_2217 | SPP_2242 | SPT_2208 | SPH_2388 | SPG_2135 | SPCG_2158 | SPN23F22240 | SPAP_2234 | SP670_2336 | SPNA45_02044 |
| *cbpC* | (360865-361143) | – | – | SP70585_0451 (405393-405722) | SPJ_0367 (354623-354952) | SPP_0418 (375155-375484) | SPT_0425 | SPH_0487 (443329-443658) | SPG_0345 (340816-341145) | (376248-376577) | SPN23F03520 (344538-344867) | SPAP_0407 | (411551-411808) | – |
| *cbpD* | SP_2201 | spr2006 | SPD_2028 | SP70585_2328 | SPJ_2227 | SPP_2254 | SPT_2219 | SPH_2399 | SPG_2147 | SPCG_2168 | SPN23F22340 | SPAP_2244 | SP670_2348 | SPNA45_02054 |
| *cbpE* | SP_0930 | spr0831 | SPD_0821 | SP70585_0969 | SPJ_0870 | SPP_0937 | SPT_1269 | SPH_1039 | SPG_0856 | SPCG_0905 | SPN23F08530 | SPAP_0962 | SP670_1390 | SPNA45_01231 |
| *cbpF* | SP_0377 | spr0337 | SPD_0345 | SP70585_0448 | SPJ_0363 | SPP_0415 | SPT_0420 | SPH_0484 | SPG_0343 | – | – | SPAP_0402 | SP670_0445 | SPNA45_01670 (1701163-1700261) |
| *cbpG* | SP_0390 | spr0350 (350513-351372) | SPD_0356 (358042-358901) | SP70585_0462 | SPJ_0378 | SPP_0430 | SPT_0436 | SPH_0499 | SPG_0356 | SPCG_0387 | SPN23F03640 | SPAP_0417 | SP670_0461 | SPNA45_01660 |
| *cbpI* | SP_0069 | – | – | – | – | – | – | – | SPG_0072 | – | – | SPAP_0116 | – | – |
| *cbpJ* | SP_0378 | – | – | – | SPJ_0364 | – | SPT_0421 | – | – | SPCG_0373 | SPN23F03490 | – | SP670_0446 | SPNA45_01670 (1700243-1699248) |
| *cbpK* | SP_0391 | spr0351 | SPD_0357 | SP70585_0464 | SPJ_0379 | – | – | SPH_0501 | SPG_0357-0358 | SPCG_0388-0389 | SPN23F03650 | – | SP670_0462 | SPNA45_01659 |
| *cbpL* | SP_0667 | spr0583 | SPD_0579 | SP70585_0726 | SPJ_0618 | SPP_0688 | SPT_0691 | SPH_0762 | SPG_0608 | SPCG_0623 | SPN23F06030 | SPAP_0656 | SP670_0726 | SPNA45_01008 |
| *cbpM* | SP_1417 | spr1274 | SPD_1248 | SP70585_1456 | SPJ_1316 | SPP_1436 | SPT_0856 | SPH_1548 | SPG_1358 | SPCG_1405 | SPN23F13820 | SPAP_1448 | SP670_0880 | SPNA45_00788 |
| *lytA* | SP_1937 | spr1754 | SPD_1737 | SP70585_2016 | SPJ_1932 | SPP_1966 | SPT_1895 | SPH_2087 | SPG_1847 | SPCG_1911 | SPN23F19600 | SPAP_1957 | SP670_2024 | SPNA45_00060 |
| *lytB* | SP_0965 | spr0867 | SPD_0853 | SP70585_1005 | SPJ_0906 | SPP_0971 | SPT_1238 | SPH_1067 | SPG_0889 | SPCG_0941 | SPN23F08900 | SPAP_0997 | SP670_1354 | SPNA45_01263 |
| *lytC* | SP_1573 | spr1431 | SPD_1403 | SP70585_1614 | SPJ_1479 | SPP_1596 | SPT_1513 | SPH_1687 | SPG_1499 | SPCG_1559 | SPN23F15900 | SPAP_1594 | SP670_1658 | SPNA45_00649 |
| *pcpA* | SP_2136 | spr1945 | SPD_1965 | SP70585_2262 | SPJ_2161 | – | SPT_2148 | SPH_2328 | SPG_2074 | SPCG_2105 | SPN23F21690 | SPAP_2186 | SP670_2281 | SPNA45_00072 |
| *pspA* | SP_0117 | spr0121 | SPD_0126 | SP70585_0197 | SPJ_0148 | SPP_0185 | SPT_0163 | SPH_0232 | SPG_0121 | SPCG_0120 | SPN23F01290 | SPAP_0173 | SP670_0200 | SPNA45_01912 |

| | TCH8431/19A | INV200 | OXC141 | INV104 | SPN034156 | SPN034183 | SPN994038 | SPN994039 | ST556 | gamPNI0373 | NT_110_58 | A66 | NCTC7465 | A026 |
|---|---|---|---|---|---|---|---|---|---|---|---|---|---|---|
| *cbpA* | HMPREF0837_10201 | – | – | INV104_18920 | SPN034156_10130 | – | – | – | MYY_2112 | HMPREF1038_02198 | – | – | – | T308_10525 |
| *cbpC* | HMPREF0837_10678 | SPNINV200_03420 | SPNOXC03770 | INV104_03270 (366826-367155) | SPN034156_14330 | SPN034183_03830 | SPN994038_03710 | SPN994039_03720 | MYY_02500 | HMPREF1038_00431 (383697-384026) | SpnNT_00359 | A66_00363 (347063-347392) | ERS445053_00114 | T308_01890 |
| *cbpD* | HMPREF0837_10214 | SPNINV200_20100 | SPNOXC19430 | INV104_19020 | SPN034156_10230 | SPN034183_19450 | SPN994038_19350 | SPN994039_19360 | MYY_2124 | HMPREF1038_02209 | SpnNT_02272 | A66_02002 | ERS445053_00568 | T308_10590 |
| *cbpE* | HMPREF0837_11555 | SPNINV200_08540 | SPNOXC08350 | INV104_07940 | SPN034156_18810 | SPN034183_08330 | SPN994038_08220 | SPN994039_08230 | MYY_1270 | HMPREF1038_00950 | SpnNT_01378 | A66_00846-00848 | ERS445053_01801 | T308_05945 |
| *cbpF* | HMPREF0837_10674 | SPNINV200_03380 | SPNOXC03730 | INV104_03240 | SPN034156_14290 | SPN034183_03790 | SPN994038_03670 | SPN994039_03680 | MYY_0454 | HMPREF1038_00428 | SpnNT_00354 | A66_00359 | ERS445053_00118 | T308_01870 |
| *cbpG* | HMPREF0837_10690 | SPNINV200_03540 | SPNOXC03870 | INV104_03370 | SPN034156_14420 | SPN034183_03920 (383887-384516) | SPN994038_03800 (373464-374093) | SPN994039_03810 (373464-374093) | MYY_0470 | HMPREF1038_00443 | SpnNT_00368 | A66_00374 | ERS445053_00104-00105 | T308_01940 |
| *cbpI* | – | – | – | – | – | – | – | – | – | – | SpnNT_00093 | – | – | – |
| *cbpJ* | HMPREF0837_10675 | SPNINV200_03390 | SPNOXC03740 | – | SPN034156_14300 | SPN034183_03800 | SPN994038_03680 | SPN994039_03690 | MYY_0455 | – | SpnNT_00355 | A66_00360 | – | T308_01875 |
| *cbpK* | – | SPNINV200_03550 | – | INV104_03380 | – | – | – | – | – | – | – | A66_00376 | – | – |
| *cbpL* | HMPREF0837_10959 | SPNINV200_05890 | SPNOXC06120 | INV104_05580 | SPN034156_16610 | SPN034183_06130 | SPN994038_06020 | SPN994039_06030 | MYY_0712 | HMPREF1038_00693 | SpnNT_00659 | A66_00605 | ERS445053_02041-02042 | T308_03150 |
| *cbpM* | HMPREF0837_11129 | SPNINV200_12620 | SPNOXC12490 | INV104_12100 | SPN034156_03360 | SPN034183_12470 | SPN994038_12360 (1258836-1257709) | SPN994039_12370 | MYY_0872 | HMPREF1038_01404 | SpnNT_00846 | A66_01258 | ERS445053_01356 | T308_03935 |
| *lytA* | HMPREF0837_12181 | SPNINV200_17580 | SPNOXC17060 | INV104_16700 | SPN034156_07870 | SPN034183_17100 | SPN994038_16990 | SPN994039_17000 | MYY_1846 | HMPREF1038_01928 | SpnNT_01991 | A66_01741 | ERS445053_00839 | T308_08980 |
| *lytB* | HMPREF0837_11523 | SPNINV200_08860 | SPNOXC08670 | INV104_08260 | SPN034156_19090 | SPN034183_08650 | SPN994038_08540 | SPN994039_08550 | MYY_1239 | HMPREF1038_00985 | SpnNT_01304 | A66_00883 | ERS445053_01767 | T308_05795 |
| *lytC* | HMPREF0837_11809 | SPNINV200_14100 | SPNOXC13820 | INV104_13450 | SPN034156_04700 | SPN034183_13810 | SPN994038_13700 | SPN994039_13710 | MYY_1506 | HMPREF1038_01558 | SpnNT_01649 | A66_01408-01409 | ERS445053_01211 | T308_07170 |
| *pcpA* | HMPREF0837_10138 | SPNINV200_19490 | – | INV104_18450 | – | – | – | – | MYY_2056 | – | – | A66_02052 | – | T308_10240 |
| *pspA* | HMPREF0837_10423 | SPNINV200_01170 | SPNOXC01620 | INV104_00930 | SPN034156_12190 | SPN034183_01680 | SPN994038_01560 | SPN994039_01560 | MYY_0199 | HMPREF1038_00189 | SpnNT_00143 | A66_00134 | ERS445053_00357 | T308_00570 |

Numbers in parentheses indicate regions in each whole genome sequence.

**Supplementary Table 2.** Prediction of ORF structures

| | **Promoter** | **−10 BOX** | **−35 BOX** | **TF-binding site** | **ORF** |
|---|---|---|---|---|---|
| SPNA45 *cbpF* | + | + | + | + | + |
| SPNA45 *cbpJ* | + | + | + | + | + |
| T4 *cbpC* | −** | + | + | + | + |
| 70585 *cbpC* | +* | + | + | + | + |
| JJA *cbpC* | +* | + | + | + | + |
| P1031 *cbpC* | +* | + | + | + | + |
| 19A-6 *cbpC* | +* | + | + | + | + |
| G54 *cbpC* | +* | + | + | + | + |
| CGSP14 *cbpC* | −** | + | + | + | + |
| ATCC 700669 *cbpC* | −** | + | + | + | + |
| 670-6B *cbpC* | −** | + | + | + | + |
| INV104 *cbpC* | +* | + | + | + | + |
| gamPNI0373 *cbpC* | +* | + | + | + | + |
| A66 *cbpC* | +* | + | + | + | + |
| R6 *cbpG* | −** | + | + | + | − |
| D39 *cbpG* | −** | + | + | + | − |
| SPN034183 *cbpG* | −** | + | + | + | + |
| SPN994038 *cbpG* | −** | + | + | + | + |
| SPN994039 *cbpG* | −** | + | + | + | + |
| SPN994038 *cbpM* | − | − | − | − | + |

*Predicted promoter position is the first nucleotide of the start codon.

**Predicted promoter position is downstream of the start codon.

**Supplementary Table 3.** Evolutionary analyses of selected *cbpC* and *cbpL* sequences

| Genes | Number of sequences | dN/dS | Coverage of comparable codons to the whole protein in TIGR4 | Codons evolving under positive selection | Codons evolving under purifying selection | % of codons under purifying selection relative to total codons |
|---|---|---|---|---|---|---|
| *cbpC* | 12 | 0.462 | 74.194% (69/93) | 0% (0/69) | 2.899% (2/69) | 2.151% |
| *cbpL* | 17 | 0.220 | 33.634% (112/333) | 0% (0/112) | 8.929% (10/112) | 3.003% |

Evolutionary analysis was performed by Bayesian inference of selected *cbpC and cbpL* sequences with the two-rate fixed-effect likelihood function in HyPhy software package. The dN/dS means ratio represents non-synonymous to synonymous changes in analysed genes. Individual codons with a statistically significant signature were also calculated and are expressed as a percentage of the total number of codons included in the analysis.

**Supplementary Table 4.** MICs and MBCs for penicillin G

| *S. pneumoniae* | Penicillin G (µg/mL) | |
|---|---|---|
| | MIC | MBC |
| WT | < 0.125 | < 0.125 |
| Δ*cbpL* | < 0.125 | < 0.125 |
| Δ*cbpJ* | < 0.125 | < 0.125 |

**Supplementary Table 5.** Primers used in this study

| Primers | Sequence (5' to 3') |
|---|---|
| T4cbpJKOuF | atcataaagcaatctattggcataa |
| T4cbpJKOuR | tattcaaatatatccctatataccctccaatattaaatcc |
| T4cbpJKOaF | ttggagggtatatagggatatatttgaatacatacgaaca |
| T4cbpJKOaR | ataagcgccctgtcatcaatttttttataattttttaat |
| T4cbpJKOdF | ttataaaaaaattgatgacagggcgcttataattatatta |
| T4cbpJKOdR | aacactctgacttttgtcattgcct |
| T4cbpLKOuF | ttgagcctaggagaacaagagaag |
| T4cbpLKOuR | tattcaaatatatccttctttctcccataattatgatatc |
| T4cbpLKOaF | ttatgggagaaagaaggatatatttgaatacatacgaaca |
| T4cbpLKOaR | gcctaccagtatctgtcaatttttttataattttttaat |
| T4cbpLKOdF | ttataaaaaaattgacagatactggtaggcgaaaaaattc |
| T4cbpLKOdR | aactttttgaacaccttgtagaagg |
| cbpL(qPCR)F | catccatttttcgagctgtaagg |
| cbpL(qPCR)R | tctccctcgtgaatatccgaat |
| cbpJ (qPCR) F | gatggcagtttgtccaagaaaa |
| cbpJ (qPCR) R | actcgccagtaggtttctttgag |
| 16s_rRNAF | tgtagcggtgaaatgcgtagata |
| 16s_rRNAR | caagccagagagccgcttt |
| lytA(qPCR)F | gctggaattaaaacgcacgagta |
| lytA(qPCR)R | ggtcaacgtggtctgagtggtt |
| pQEcbpJOPTiF | caccatcaccatcacgacgattccgaaggttggcagtttg |
| pQEcbpJOPTiR | aagctcagctaattaacgaacccactcgccattatagttc |
| pQEcbpJOPTvF | ggcgagtgggttcgttaattagctgagcttggactcctgt |
| pQEcbpJOPTvR | accttcggaatcgtcgtgatggtgatggtgatgcgatcct |
